# Intracellular pathogen targeting by IL32 elicits cell-autonomous immunity

**DOI:** 10.64898/2026.07.21.739902

**Authors:** Jeffrey R Reitano, Stephen C Walsh, Mary S Dickinson, Emma A Johnston, María E Cortina, Haley Adcox, Neal M Alto, Isabelle Derre, Kevin Hybiske, Robert Suchland, So Young Kim, Jörn Coers

## Abstract

Interferon-γ safeguards humans against intracellular pathogens, yet how most interferon-stimulated genes protect host cells, and how human-adapted pathogens evade these defenses is unclear^1,2^. Here, we discover a potent immune surveillance and effector circuit executed by an intracellularly acting cytokine, IL32, that targets and restricts phylogenetically distinct vacuolar pathogens, including the bacterium *Chlamydia* and the microsporidian *Encephalitozoon*. Quantitative proteomics coupled to a tailored CRISPR screen, uncovered components of the cysteine/Arg N-degron pathway^3^ that modify IL32 through oxidation-dependent arginylation, thereby enabling the recruitment of the autophagy machinery to pathogen-containing vacuoles. A forward genetics screen in *Chlamydia trachomatis*, the leading cause of sexually transmitted bacterial infection, identified the secreted virulence factor IncS as an evasion factor that blocks IL32 targeting and shields this human pathogen from xenophagy. These findings establish an IL32-dependent intracellular sensing mechanism linking IFNγ signaling to N-degron–mediated xenophagy, revealing a broadly relevant axis of human host–pathogen conflict.

## IL32 targets and restricts two phylogenetically distinct intravacuolar pathogens

*Chlamydia muridarum* (*Cm*) is a close relative of the human pathogen *Chlamydia trachomatis* (*Ct*), sharing a nearly syntenic genome with only ∼10% difference in orthologous genes^4^. Despite this similarity, as a rodent-adapted pathogen, *Cm* lacks many of *Ct*’s human-specific immune evasion strategies, making *Cm* a useful tool for the study of human anti-*Chlamydia* immunity^5^. Both *Ct* and *Cm* productively infect cultured human epithelial cells, residing in membrane-bound vacuoles termed inclusions^6^. However, when such cells are primed with interferon-gamma (IFNγ), *Cm* growth is restricted whereas *Ct* evades this restriction^7,8^ (Fig. 1a). We inactivated both known pathways of human IFNγ-mediated cell autonomous defense against *Cm* by genetically deleting the ubiquitin E3 ligase RNF213 and supplementing excess tryptophan to reverse the effects of the tryptophan-degrading enzyme indole-dioxygenase (IDO)^7,9^. Nonetheless, we found significant restriction of *Cm* that was independent of both pathways (Fig. 1b), indicating the existence of at least one additional human anti-*Chlamydia* interferon-stimulated gene (ISG) and a corresponding *Ct* evasion mechanism (Fig. 1c).

**Figure 1.**
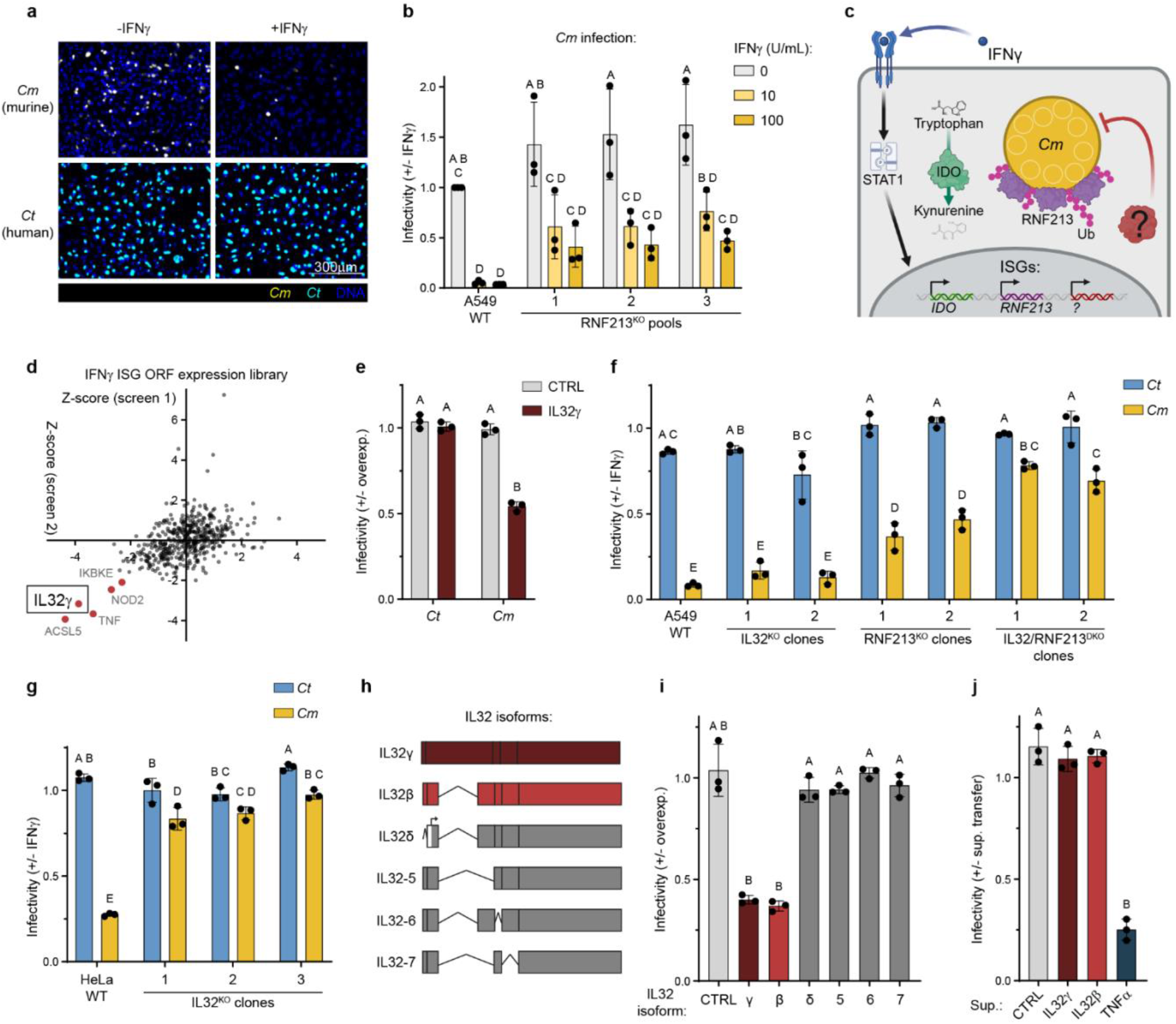
ISG overexpression screen identifies IL32 as a novel anti-*Chlamydia* host factor. Representative images of HeLa cells (nuclei, blue) primed overnight with 0 or 100U/mL IFNγ and subsequently infected with *Cm* (yellow) or *Ct* (cyan) for 24 hours. **b.** Relative infectivity of A549 cells (WT or RNF213^KO^) primed overnight with 0, 10, or 100U/mL IFNγ, treated with 100µg/mL L-Trp, and subsequently infected with *Cm* at a multiplicity of infection (MOI) of 2 for 24 hours. Three independent repeats were quantified by high-content imaging and are shown as mean ± S.D. **c.** Diagram depicting IFNγ-induced anti-*Chlamydia* cell-autonomous immune pathways. **d.** Dot plot showing results of two independent ISG overexpression screens. 470 ISGs were individually overexpressed in A549 cells and cells were infected with *Cm* for 24 hours. Z-scores of *Cm* inclusions per host nuclei are depicted, and genes with a Z-score <-2 (p<0.05) in both repeats were identified as “hits”. **e.** IL32 or a firefly luciferase (fLuc) control were overexpressed in A549s for 24 hours before infection. Relative infectivity of *Ct* and *Cm* are shown. **f.** Relative infectivity of *Ct* and *Cm* in IFNγ-primed A549s, comparing parental cells (WT) to IL32^KO^, RNF213^KO^, and IL32/RNF213^DKO^ clones. **g.** Relative infectivity of *Ct* and *Cm* in IFNγ-primed HeLas, comparing a wild-type pool to IL32^KO^ clones. **h.** Diagram of six IL32 isoforms investigated in this study. **i.** Relative infectivity of *Cm* in A549 cells expressing six FLAG-tagged isoforms of IL32, or a TagRFP control. **j.** Supernatant was taken from A549 cells expressing IL32γ, IL32β, TNFα, or a fLuc control and transferred onto fresh A549 cells overnight. Cells were subsequently infected with *Cm* and infectivity was measured relative to a “no transfer” control. Data in **b**, **e**, **f**, and **g** were analyzed with a two-way ANOVA with Tukey’s multiple comparisons test. Data in **i** and **j** were analyzed with a Brown-Forsythe and Welch one-way ANOVA with Dunnett’s T3 multiple comparisons test. Any groups that do not share a letter are statistically different; p<0.05.

To identify this predicted ISG, we used a lentiviral expression library^10^ to individually express 470 human IFNγ-responsive ISGs in A549 cells and measured restriction of *Cm*. For this screen, and all subsequent restriction assays, we included 100µg/mL L-Tryptophan to nullify IDO. This screen yielded a single validated hit not previously linked to host defense against *Chlamydia*: the orphan cytokine interleukin-32 (IL32) (Fig. 1d; Supplemental Table 1). Ectopic expression of IL32 alone in A549 cells was sufficient to restrict *Cm* but not *Ct*, and deletion of IL32 in RNF213^KO^ A549 cells resulted in a further loss of *Cm* restriction (Fig. 1e,f; S1a,b). In HeLa cells, IL32 deletion alone nearly abolished all IFNγ-mediated restriction of *Cm* (Fig. 1g; S1c,d). Collectively, these results establish IL32 as a novel anti-*Chlamydia* ISG. Since IL32 has been extensively studied as a cytokine^11^, we initially hypothesized an extracellular, autocrine role against *Cm*. Among all IL32 splice isoforms, only the full-length IL32γ has a potential secretion signal sequence^12^, yet both IL32γ and IL32β restricted *Cm* (Fig. 1h,i). Additionally, IL32 was predominantly intracellular in HeLa and A549 cells, and the transfer cell supernatants harvested from IL32 expressing cells failed to confer protection against *Cm* (Fig. 1j; Fig. S1e). These findings indicate that IL32 restricts *Cm* via a novel, intracellular mechanism.

To define this unexpected activity of IL32, we used immunofluorescence to map its subcellular localization. Endogenous IL32 selectively decorated inclusion membranes containing *Cm*, but not *Ct*, across HeLa, A549, and primary HCerEpiC epithelial cells, and HFF-1 fibroblasts (Fig. 2a-d; S2a-c). All six IL32 splice isoforms tested efficiently targeted *Cm* inclusions (Fig. 2e), indicating that the determinants of inclusion binding are shared between these isoforms. In contrast, determinants of inclusion destruction are missing from the shorter δ, 5, 6, and 7 isoforms (Fig. 1i). Motivated by the possibility that IL32-mediated restriction in human cells might extend to other maladapted vacuolar pathogens, we examined the opportunistic fungal parasite *Encephalitozoon cuniculi*^13^ and found that its vacuoles were likewise targeted by endogenous IL32, and that growth of *E. cuniculi* was curtailed by IL32β overexpression (Fig. 2f–h). These findings establish the cytokine IL32 as a previously unrecognized intracellular defense protein that directly targets and eliminates diverse intravacuolar pathogens.

**Figure 2.**
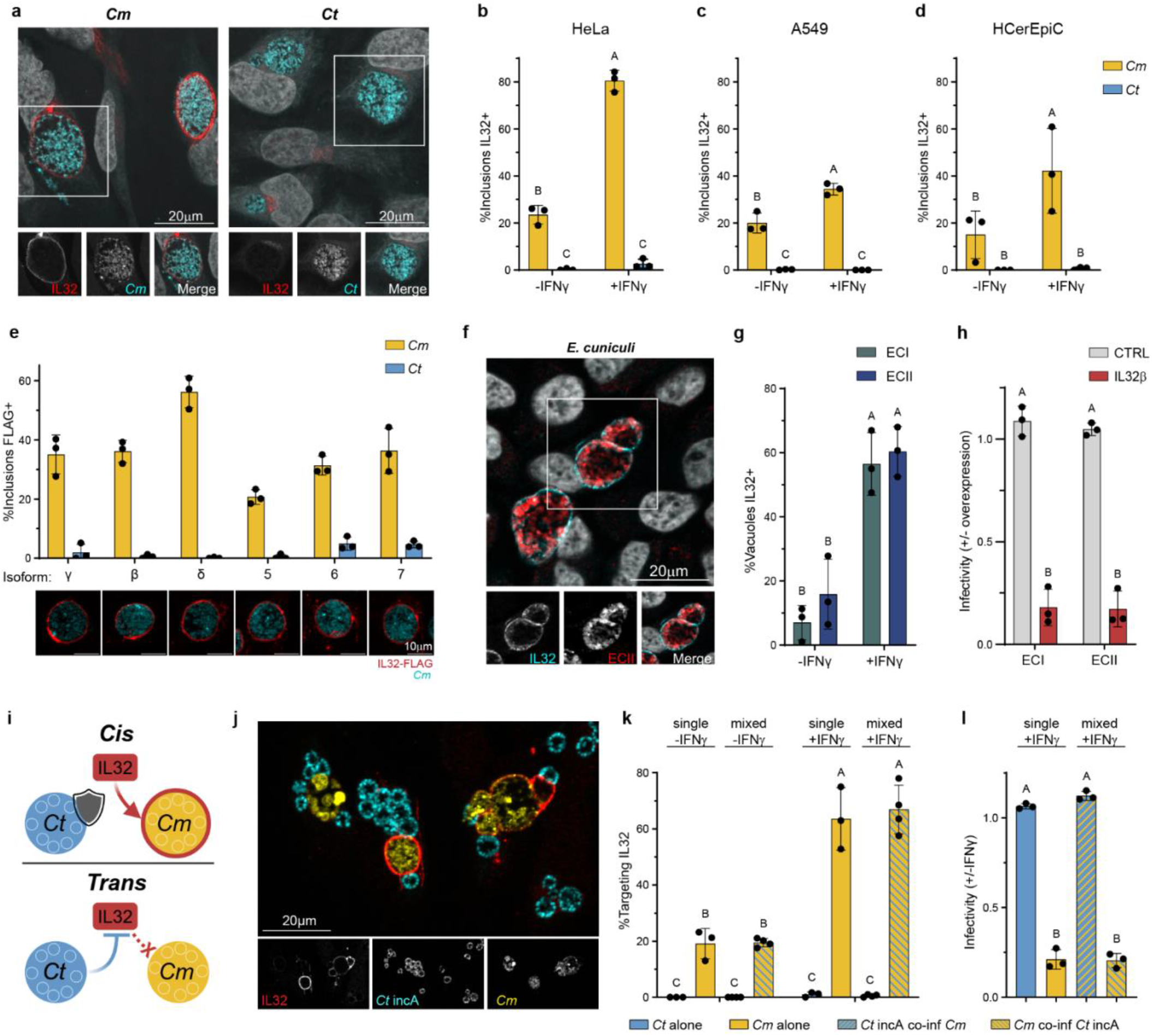
IL32 directly targets *Cm*- and *Microsporidia*-containing vacuoles. **a.** Representative image of IL32 (red) localizing to *Cm* but not *Ct* (cyan) vacuoles at 20hpi in HeLa cells. **b.** Quantification of IL32 targeting to *Cm* or *Ct* inclusions in IFNγ-primed or unprimed HeLa cells, **c.** A549 cells, and **d.** HCerEpiC primary human cervical epithelial cells. Each datapoint represents at least five fields-of-view, typically 80+ inclusions. **e.** Quantification and representative images of FLAG-tagged IL32 isoforms (red) to *Cm* inclusions (cyan) or *Ct* inclusions. **f.** Representative image of IL32 (cyan) targeting to *E. cuniculi* strain ECII (red), and **g.** quantification in HeLa cells. **h.** IL32β or a fLuc control was overexpressed in A549 cells for 24 hours, cells were infected with ECI or ECII at an MOI of 20, and infectivity was quantified with high content imaging at 50hpi. **i.** Diagram depicting *Ct* evasion of IL32 *in cis* or *in trans*. **j.** Representative image of co-infection, measuring targeting of IL32 (red) to *Cm* (yellow) and *Ct* incA^-^(cyan) in co-infected HeLas and **k.** quantification of IL32 targeting. **l.** Relative infectivity of *Ct*, *Cm*, or *Ct* incA^-^ with or without co-infection in HeLa cells. Data in **b**, **c**, **d**, **g**, **h**, **k**, and **l** were analyzed with a two-way ANOVA with Tukey’s multiple comparisons test.

## The *Ct* virulence factor IncS blocks IL32-mediated defense

We next asked how human-adapted *Ct* evades IL32 targeting and restriction. We first tested whether *Ct* blocks IL32 *in cis* (at the site of individual inclusions) or *in trans* (globally throughout the cell). To this end, we co-infected cells with *Cm* and an IncA-deficient *Ct* mutant. As IncA is required for inclusion fusion, *Cm* and *Ct* in the same cell remain in separate inclusions in the absence of IncA^14,15^ (Fig. 2i). IL32 targeting and restriction of *Cm* was unaffected by the presence of *Ct* within the same cell, demonstrating that evasion occurs *in cis* (Fig. 2j-l). Furthermore, this suggests that IL32 inclusion binding is required for its antimicrobial activity.

To map the genetic locus encoding the putative IL32 evasion factor, we leveraged a mini library of twenty chimeric *Chlamydia* strains carrying different segments of the *Ct* and *Cm* genomes^16,17^ (Fig. 3a). We measured IL32 targeting, restriction in IFNγ-primed HeLa cells, and restriction in IL32β-overexpressing A549s for each strain. Among these, RC826 and RC1323 differ at only a single locus encoding the inclusion membrane protein IncS (CTL0402) (Fig. 3a), yet RC826 was robustly targeted and restricted by IL32, whereas RC1323 was not (Fig. 3b-e). These data implicated *Ct* IncS as an IL32 evasion factor.

**Figure 3.**
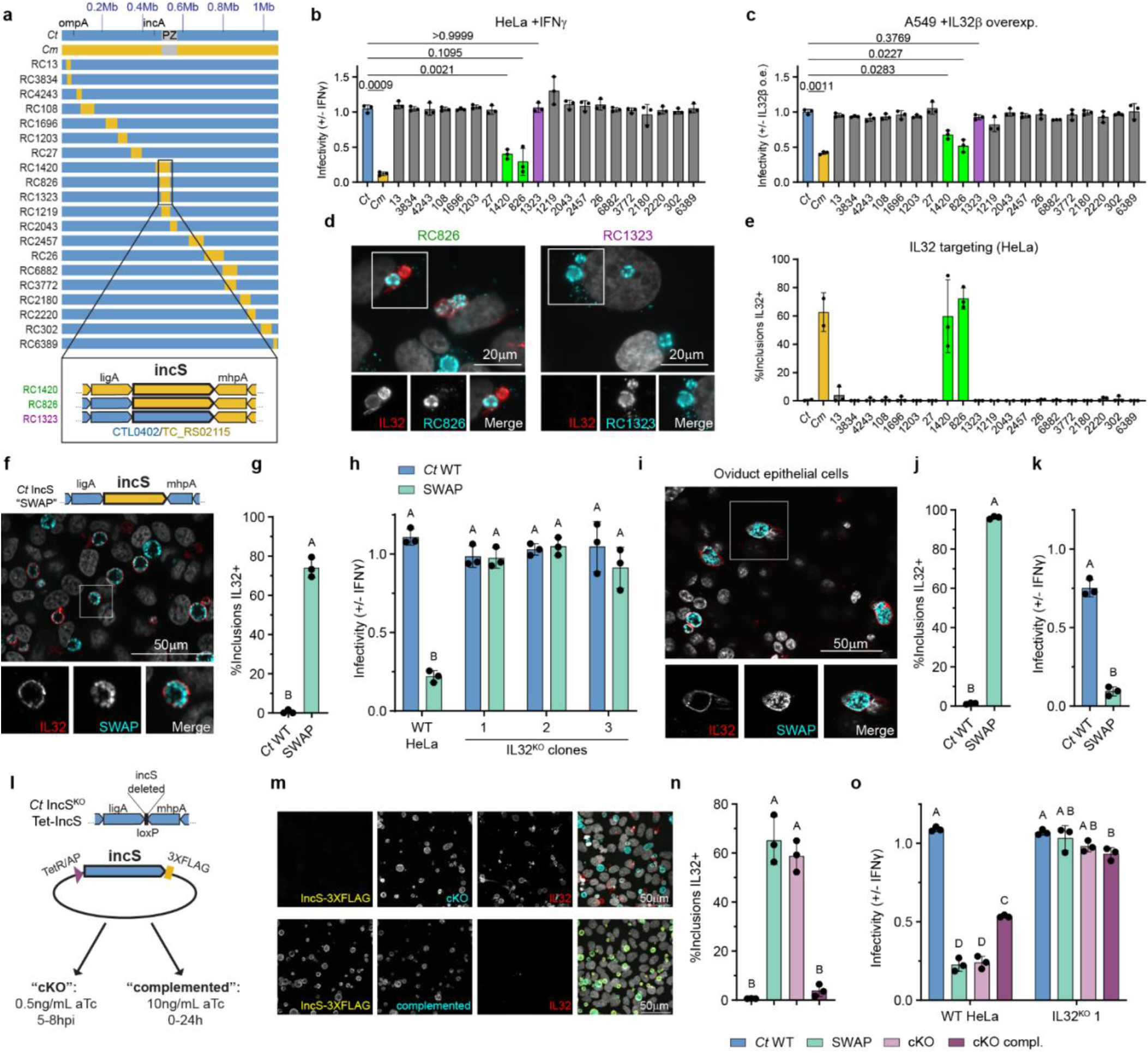
*Ct* IncS prevents IL32 inclusion targeting. **a.** Diagram of *Cm*-*Ct* chimera genomes, including a zoom in on IncS, the only locus in which RC826 and RC1323 differ. **b.** Relative infectivity of chimeras in IFNγ-primed HeLas and **c.** in IL32β-overexpressing A549s. **d.** Representative images and **e.** quantification of IL32 targeting to chimeras (n=3) compared to *Cm* and *Ct* controls (n=2). Each datapoint represent four fields-of-view in IFNγ-primed HeLa cells. **f.** Genetic diagram and representative image of IL32 targeting *Ct* SWAP in IFNγ-primed HeLas. **g.** IL32 targeting to WT *Ct* and *Ct* SWAP in IFNγ-primed HeLas. **h.** Relative infectivity of *Ct* and SWAP in IFNγ-primed HeLas (WT and IL32^KO^ clones). **i.** Representative image, **j.** IL32 targeting data, and **k.** relative infectivity of *Ct* SWAP and WT *Ct* in IFNγ-primed E6/E7-immortalized oviduct epithelial cells. **l.** Genetic diagram of *Ct* IncS^KO^ Tet-IncS strain. **m.** Representative images in IFNγ-primed HeLa cells of IL32 (red) targeting *Ct* IncS^KO^ Tet-IncS strain (cyan) under cKO conditions, but not cKO complemented conditions, where IncS (yellow) is visible. **n.** Corresponding IL32 targeting data, including *Ct* and SWAP controls. **o.** Relative infectivity of *Ct*, IncS SWAP, IncS cKO, and IncS complemented strains in WT HeLas or IL32^KO^ clone 1. Data in **b**, **c**, and **n** were analyzed with a Brown-Forsythe and Welch one-way ANOVA with Dunnett’s T3 multiple comparisons test. Data in **g**, **j**, and **k** were analyzed with an unpaired, two-tailed Welsh’s t-test. Data in **h** and **o** were analyzed with a two-way ANOVA with Tukey’s multiple comparisons test.

IncS is essential for chlamydial development between 6-8 hpi^18^, precluding generation of a conventional IncS gene deletion in *Ct*. We therefore used an IncS “SWAP” strain in which endogenous *Ct* IncS was chromosomally replaced with the *Cm* ortholog^19^ (Fig. 3f). This SWAP strain grows normally in unprimed cells lacking significant IL32 expression^19^ but we found that it was robustly targeted by IL32 and restricted when IL32 is expressed (Fig. 3f-h; S3a-c). Consistently, IFNγ-primed E6/E7-immortalized human oviduct epithelial cells^20^ efficiently IL32-decorated and restricted over 90% of IncS SWAP inclusions (Fig. 3i-k). Therefore, replacement of *Ct* IncS with *Cm* IncS transfers IL32 sensitivity to *Ct.* However, this approach did not distinguish whether *Cm* IncS lacks the IL32-evasion function of *Ct* IncS, or whether *Cm* IncS actively promotes IL32 recruitment. To differentiate between these two models, we employed the *Ct* IncS^KO^ Tet-IncS strain, which lacks chromosomal IncS but expresses *Ct* IncS from a tetracycline-inducible plasmid^18^. We compared a “complemented” condition with 10ng/mL aTc present throughout infection, yielding high *Ct* IncS expression, to a “conditional KO (cKO)” condition with 0.5ng/mL aTc present only from 5-8 hpi, which preserves bacterial viability but dramatically reduces IncS expression (Fig. 3l, m). Under cKO conditions, inclusions were targeted and restricted by IL32, whereas IncS complementation conditions markedly reduced IL32 targeting and restriction (Fig. 3m-o). Together, these data identify the inclusion protein *Ct* IncS as the singular virulence factor that enables *Ct* to evade IL32-mediated targeting and restriction *in cis*.

## IL32 activates N-degron-dependent xenophagy

We next investigated the mechanism of IL32-mediated restriction. Although IL32 has been reported to induce caspase-3-dependent cell death^21,22^, IL32 overexpression during *Cm* infection did not trigger cytotoxicity, and blockade of caspases with the pan-caspase inhibitor ZVAD did not abrogate the anti-*Chlamydia* activity of IL32 (Fig. S4a,b). IL32 has also been implicated in the induction of pro-inflammatory cytokines^23,24^, yet conditioned supernatants from IL32-expressing cells failed to confer anti-chlamydial activity (Fig. 1j), indicating that no secreted factor induced by IL32 is sufficient to account for its cell-autonomous restriction of *Chlamydia*. Moreover, IL32-mediated restriction was fully preserved in the absence of the anti-*Chlamydia* ubiquitin E3 ligase RNF213, and IL32 was not active against the RNF213-sensitive *Ct* garD:GII mutant (Fig. S4c,d). Collectively, these data together indicate that IL32 orchestrates a novel host defense program.

Because both IL32β and IL32δ localize to *Cm* but only IL32β mediates restriction (Fig. 1h, 2f), we hypothesized that IL32δ fails to engage protein partners essential for antimicrobial activity. Pursuing this hypothesis, we devised a TurboID-based proximity labeling screen^25^ to identify key IL32 interaction partners required for host defense (Fig. 4a; Supplemental Table 2). A549 cells expressing IL32β-TurboID, IL32δ-TurboID, or TurboID alone were infected with *Cm* for 4 hours; labelled with biotin for an additional 5h; and lysed for streptavidin pulldown and LC-MS/MS. To optimize the labeling of IL32β interaction partners, we used four single-cell clones expressing high levels of IL32β-TurboID (Fig. S5a) and included parallel samples treated with the proteasomal inhibitor epoxomicin to retain short-lived biotinylated proteins. Like their untagged counterparts, IL32β-TurboID and IL32δ-TurboID both target *Cm*, but only IL32β-TurboID restricts *Cm* (Fig. S5b,c).

**Figure 4.**
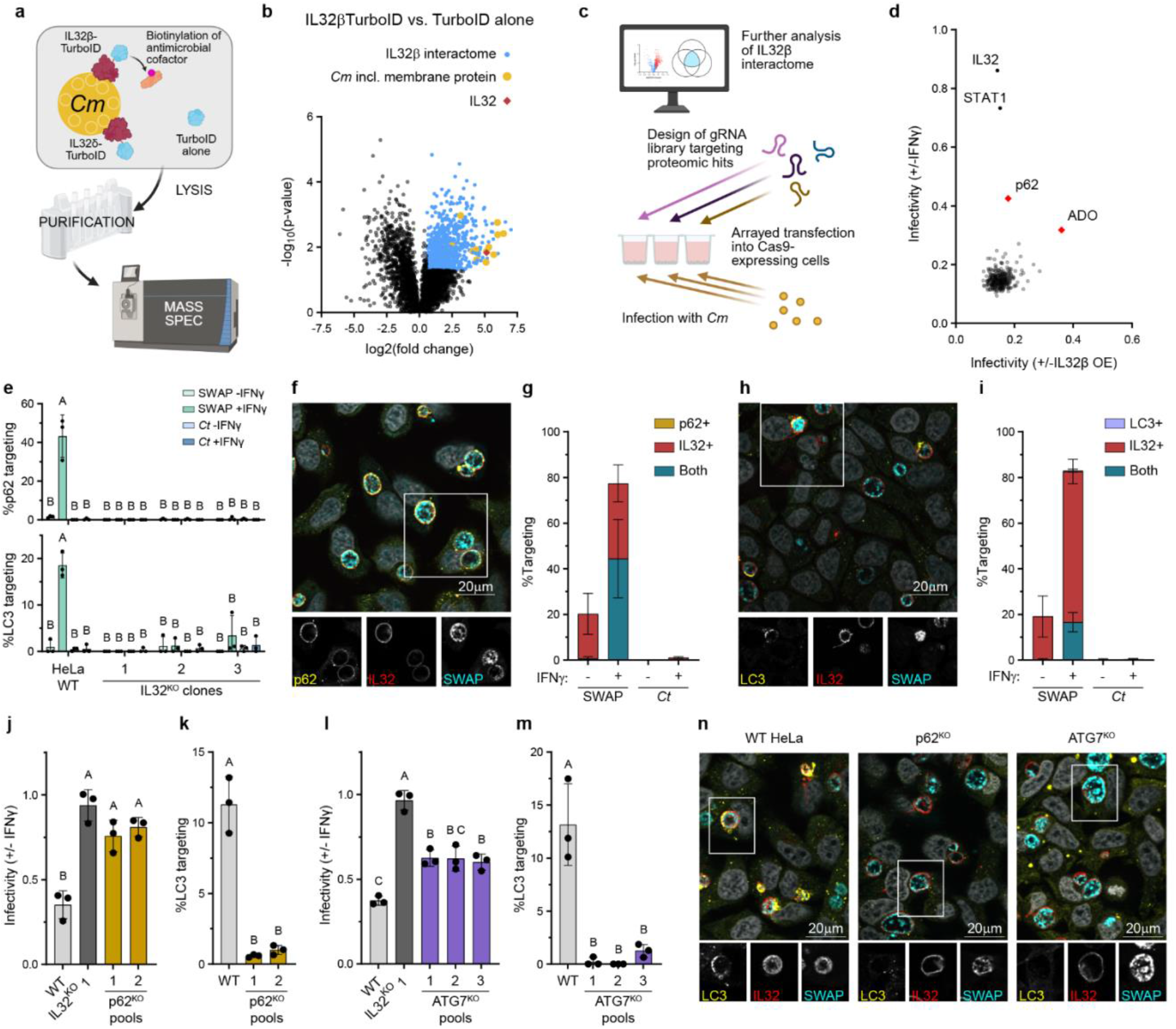
Functional screen of the IL32 interactome reveals p62-dependent xenophagy operates downstream from IL32. **a.** Diagram of TurboID-based proteomic screen to identify IL32 interactors. **b.** Volcano plot showing IL32β interactome during *Cm* infection. Blue circles are proteins with a fold-change>1.5 and p-value<0.05 in IL32β-TurboID cells compared to TurboID cells. **c.** Diagram and **d.** results from an arrayed CRISPR screen probing the function of 289 putative IL32 interacting proteins, two positive controls (IL32 and STAT1), and two negative controls (AAVS1 and scrambled). Average relative infectivity of *Cm* is depicted in IFNγ-primed HeLa cells (Y-axis, n=3) or IL32β-overexpressing A549 cells (X-axis, n=2), **e.** p62 and LC3 targeting to *Ct* SWAP in HeLa cells (WT and IL32^KO^ clones). **f.** Representative image and **g.** data showing co-localization of IL32 and p62 on *Ct* SWAP in HeLa cells. **h.** Representative image and **i.** data showing IL32 and LC3 co-localization on *Ct* SWAP in HeLa cells. **j.** Relative SWAP infectivity in p62^KO^ pool HeLa cells, with parental (WT) and IL32^KO^ controls. **k.** LC3 targeting in WT and p62^KO^ pool HeLa cells. **l.** Relative SWAP infectivity in WT, IL32^KO^, and ATG7^KO^ pool HeLa cells. **m.** LC3 targeting in WT and ATG7^KO^ pool HeLa cells. **n.** Representative images of IL32 and LC3 targeting to *Ct* SWAP in IFNγ-primed WT, p62^KO^ pool 2, and ATG7^KO^ pool 1 HeLa cells. Data in **e** were analyzed with a two-way ANOVA with Tukey’s multiple comparisons test. Data in **j-l** were analyzed with a Brown-Forsythe and Welch one-way ANOVA with Dunnett’s T3 multiple comparisons test. **m** was analyzed with a one-way ANOVA with Tukey’s multiple comparisons test.

We first defined an “IL32β interactome” as those proteins more abundant in IL32β-TurboID than in TurboID alone (p-value<0.05, Fold change>1.5) (Fig. 4b). From this set, we isolated two subsets for downstream analysis. The first, “β-specific” proteins, comprises host factors enriched with IL32β-TurboID compared to IL32δ-TurboID (p<0.05, fold-change>1.5; and fold-change>4 in the IL32βTurboID vs. TurboID comparison), representing candidates that IL32δ fails to recruit. This category contained 189 proteins. Because IL32β and IL32δ also share a subset of binding partners, some of which could be important for antibacterial activity, we next defined a “core set” of proteins that are common to both IL32 isoform interactomes. Specifically, this core set comprises 107 proteins that were significantly enriched (p<0.05, fold-change>1.5) in all three of the following comparisons: IL32β-TurboID vs. TurboID, IL32δ-TurboID vs. TurboID, and IL32β-TurboID with proteasome inhibition vs. TurboID with proteasome inhibition. Together, the IL32β-specific and core sets yielded 289 candidate interaction partners, with seven proteins shared between the two subsets, that collectively define a high-confidence IL32-associated protein network formed during *Chlamydia* infection (Fig. S5d).

We next used an arrayed gRNA library targeting each of the 289 candidate interactors to define which IL32 partners are required for pathogen restriction. Genes were individually disrupted in both IFNγ-primed HeLa-Cas9 cells and in IL32β-overexpressing A549-Cas9 cells, followed by *Cm* infection (Fig. 4c; Supplemental Table 3). Knockouts in two genes diminished IL32-mediated host defense: the cysteine-modifying enzyme 2-Aminoethanethiol Dioxygenase (ADO) and the autophagy adaptor Sequestosome-1 (p62) (Fig. 4d). Given the well-established role of p62 in selective autophagy^26^, we tested the hypothesis that IL32 recruits p62 to vulnerable *Chlamydia* inclusions to promote autophagic bacterial clearance. Consistent with this model, we observed robust, IFNγ-dependent recruitment of p62 and the autophagosome marker LC3 to SWAP but not WT *Ct* inclusions. This recruitment was IL32-dependent, as IL32 deletion eliminated nearly all p62 and LC3 targeting (Fig. 4e). Furthermore, nearly all inclusions marked by p62 or LC3 were also decorated with IL32 (Fig. 4f-i), supporting a model in which IL32 directly recruits these factors. Both p62 and downstream autophagy components are required for efficient control of the SWAP strain. LC3 targeting and restriction of SWAP were lost in p62-deficient cells (Fig. 4j,k; S6a) and knockouts in the essential autophagy components ATG7 and ATG5 similarly impaired LC3 recruitment and full restriction of SWAP (Fig. 4l,m; S6b-e). In line with p62 and autophagy as downstream effectors of IL32, we found IL32 targeting was unchanged in p62, ATG7, or ATG5 knockouts (Fig. 4n; S6f-h). Autophagy is unlikely to be the sole IL32 effector pathway, because SWAP is still partially restricted by IL32 in autophagy-deficient HeLa cells (Fig. 4l; S6d).

Our second hit, ADO, oxidizes N-terminal cysteines, which are subsequently arginylated by arginyltransferase-1 (ATE1) to generate N-terminal “degrons” on a variety of proteins^27,28^. Canonically, N-degrons target proteins for proteasomal degradation^3^, but it remains largely unexplored whether this powerful quality-control pathway is utilized in cell-autonomous immunity. We found that ADO contributed to bacterial restriction in a context-dependent manner: loss of ADO did not impair restriction of SWAP in IFNγ-primed HeLa cells (Fig. S7a,b), but completely abolished restriction of SWAP by IL32β overexpression (Fig. S7c,d). Whereas ADO can be functionally redundant with other oxidation pathways^29^, arginylation by ATE1 represents an essential enzymatic step in the Cys/N-degron pathway^3^. Consistent with this, ATE1 was universally essential for SWAP restriction. Pharmacological inhibition of ATE1 with tannic acid disrupted restriction in IL32-overexpressing A549 cells (Fig. 5a; S7e) and genetic deletion of ATE1 in IFNγ-primed HeLa cells similarly blocked restriction of SWAP (Fig. 5b; S7f). IL32 targeting to inclusions was unchanged in ATE1-deficient cells, indicating that N-degron formation acts downstream of IL32 recruitment (Fig. 5c,d; S7g). These findings reveal ATE1-dependent N-degron signaling as a critical and previously unrecognized arm of cell-autonomous antibacterial immunity.

**Figure 5.**
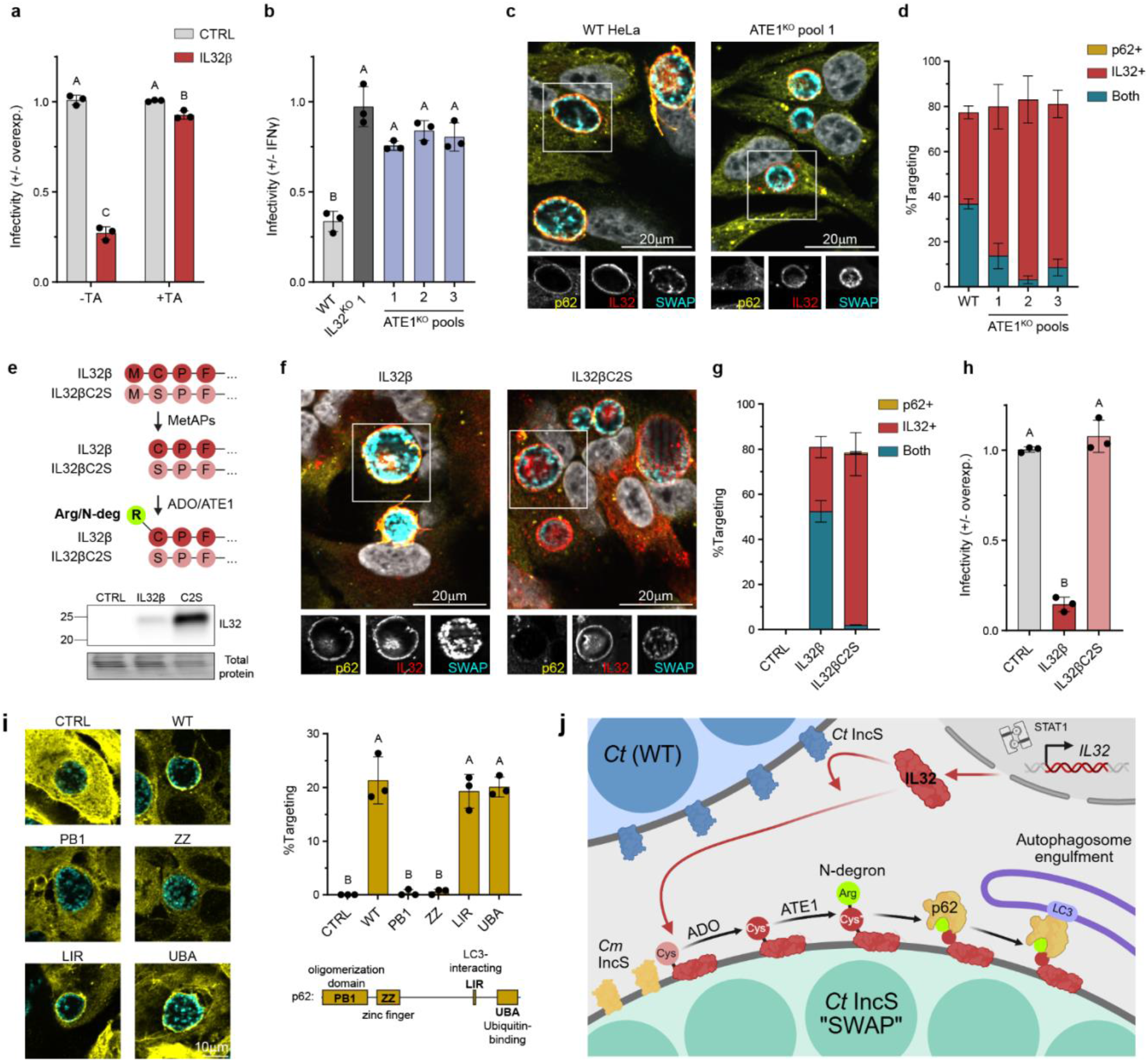
The N-degron pathway is required for IL32-mediated restriction. **a.** Relative SWAP infectivity in fLuc or IL32β-overexpressing A549s treated with 0 or 80µM tannic acid (TA). **b.** Relative SWAP infectivity in IFNγ-primed WT, IL32^KO^, and ATE1^KO^ HeLa cells. **c.** Representative images and **d.** quantification of IL32 and p62 targeting SWAP in IFNγ-primed WT and ATE1^KO^ HeLa cells. **e.** Diagram and Western blot depicting IL32β and the IL32βC2S mutant and their expression levels. **f.** representative images of p62 targeting (yellow) to SWAP (cyan) in A549 IL32^KO^ clone 1 overexpressing either IL32β or IL32βC2S mutant (red), and **g.** quantification of targeting. **h.** Relative SWAP infectivity in A549 IL32^KO^ clone 1 overexpressing IL32β, IL32βC2S, or a no vector control. **i.** Representative images, quantification, and diagram of FLAG-TurboID-tagged p62 constructs targeting to SWAP in IFNγ-primed p62^KO^ HeLa clone 1. **j.** Graphical abstract depicting IL32 targeting the *Chlamydia* inclusion, being modified by N-degron enzymes, recruiting p62, and degrading the inclusion through autophagy. Data in **a** were analyzed with a two-way ANOVA with Tukey’s multiple comparisons test. Data in **b** and **h** were analyzed with a Brown-Forsythe and Welch one-way ANOVA with Dunnett’s T3 multiple comparisons test. Data in **i** were analyzed with a one-way ANOVA with Tukey’s multiple comparison’s test.

Beyond proteasomal degradation, N-degrons can recruit p62 to drive selective autophagy of diverse intracellular structures including the endoplasmic reticulum, peroxisomes, and lipid droplets^30–32^, but they have not yet been found to target invading pathogens. Since IL32 contains an N-terminal cysteine and is itself a known substrate of ADO^27^, we hypothesized that the N-degron of IL32 could directly recruit p62 to *Chlamydia* inclusions to initiate antimicrobial autophagy. In support of this model, ATE1-deficient cells showed a marked defect in p62 recruitment compared to WT controls (Fig. 5c,d; S7h). To test whether this reflected direct modification of IL32, we manipulated the second residue of IL32β. Substitution of cysteine 2 with the small, nonpolar alanine to prevent N-degron formation also abolished most inclusion targeting. However, substitution with serine, which also blocks N-degron formation but maintains the size and polarity of cysteine, preserved inclusion targeting (Fig. S7i). Strikingly, the IL32βC2S mutant, despite robust targeting of SWAP inclusions, failed to recruit p62 or restrict *Chlamydia* (Fig. 5f-h; S7j). This indicates that N-degron formation on IL32 is specifically required to license downstream effector recruitment and subsequent pathogen killing.

In some contexts, p62 recognizes N-degrons using its zinc finger (ZZ) domain^30^, and recent work shows that p62-IL32 interactions during oxidative stress depend on this module^33^. We therefore mapped the p62 regions necessary for inclusion targeting. The zinc finger (ZZ) and polymerization (PB1) domains were essential for full recruitment to SWAP inclusions, while the ubiquitin-binding (UBA) and LC3-interacting (LIR) regions were dispensable (Fig. 5i). Together, these findings reveal that the intracellular cytokine IL32 acquires an ATE1-dependent N-degron to recruit p62 and autophagy machinery for pathogen restriction, thus establishing targeted N-degron signaling as a new principle of cell-autonomous antimicrobial immunity (Fig. 5j).

## Discussion

IL32 has been implicated in a variety of human diseases including rheumatoid arthritis and cancer^11,34^, but the basis of its evolutionary selection in humans is still poorly understood. Despite its annotation as a cytokine, it shares no homology with other cytokines, is predominantly intracellular in most contexts, and lacks a defined IL32-specific receptor^11^. Our work uncovers a previously unrecognized function for IL32 as an intracellular executioner of cell-autonomous immunity in human cells, wherein IL32 localizes to pathogen-containing vacuoles, acquires an N-degron at its N-terminal cysteine, and uses this N-degron to recruit p62 and the autophagy machinery for pathogen destruction. The IL32 pathway targets and destroys at least two evolutionarily distant intravacuolar pathogens, *C. muridarum* and *E. cuniculi*. IL32 has been shown to defend against an even broader spectrum of pathogens^35^, suggesting that it this IL32/N-degron axis may define a generalizable strategy for intracellular antimicrobial defense in humans. IL32 is encoded in humans and most other mammals but conspicuously absent in mice^11^. In this light, the IL32 susceptibility of *Cm*, which evolved in IL32-deficient murine hosts, likely exemplifies how IL32 creates a host-specific evolutionary barrier that invading pathogens must navigate, potentially contributing to host tropism and limiting zoonotic infections in humans.

The N-degron pathway is well established as an immune regulator in plants, where it primarily controls transcriptional programs rather than direct pathogen destruction^36,37^. In mammalian cells, N-degrons have been shown to recruit autophagy machinery for selective clearance of sterile intracellular structures such as the endoplasmic reticulum, peroxisomes, and lipid droplets^30–32^. In the context of infection, an N-degron within NLRP1 is crucial for the inflammasome response to anthrax lethal toxin and other virulence factors^38–40^, but N-degron signaling had not been linked to direct killing of intracellular pathogens. Here we have shown that IL32 acquires a Cys/N-degron to recruit p62 and autophagy machinery to pathogen-containing vacuoles, thus expanding the reach of N-degron biology from quality control of self-components and inflammasome licensing to a novel role in targeting foreign invaders. These results establish targeted N-degron signaling as a central mechanism of human cell-autonomous antimicrobial immunity and position IL32 as a cytokine-encoded “address label” that directs autophagic effectors to sites of infection.

*Ct* is highly adept at evading cell-autonomous immune responses in humans, and the immune system more broadly, to cause chronic infection and resulting disease^5,41^. We have identified the *Ct* inclusion membrane protein IncS as a dedicated virulence factor that enables *Ct* to evade IL32-dependent targeting and degradation in human epithelial cells. IncS thus adds to the expanding repertoire of *Ct* immune evasion strategies, which include a partial trp operon and persistence program to withstand IDO-mediated tryptophan starvation, and expression of the effector GarD to avoid RNF213-dependent inclusion ubiquitylation and destruction^9,42,43^. The existence of a specific inclusion protein that neutralizes IL32/N-degron signaling underscores the selective pressure exerted by this pathway on professional human pathogens. More broadly, these findings highlight the IL32 degradation axis as a critical barrier that successful intracellular pathogens must overcome and suggest that evasion of host-specific N-degron degradation pathways may be a recurring theme in the evolution of host tropism.

---

**Figure S1.**
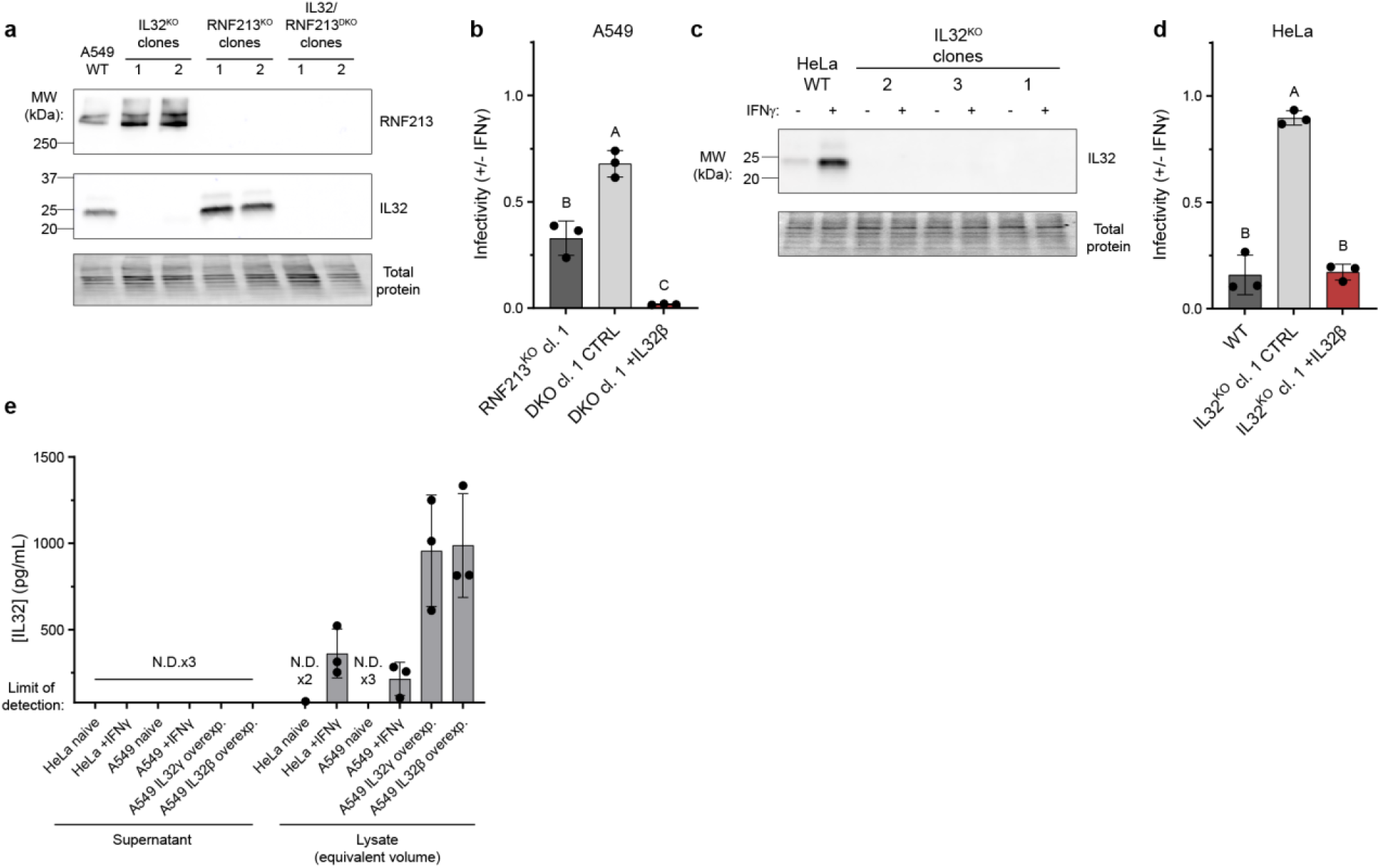
Total and supernatant IL32 protein levels and functional complementation of IL32^KO^ cells. **a.** Western blot measuring RNF213, IL32, and total protein (using UV-treated stain-free gels) on lysates of IFNγ-treated A549 cells. A wild-type pool or knockout clones for IL32, RNF213, or both genes were used. **b.** A549 IL32/RNF213^DKO^ clone 1 was complemented with pInducer20 containing IL32β or a fLuc control. Respective genes were overexpressed at the time of IFNγ stimulation. Relative *Cm* infectivity is shown. **c.** Western blot measuring IL32 and total protein on lysates from wild-type HeLas or IL32 knockout clones. **d.** HeLa IL32^KO^ clone 1 was complemented with pInducer20 containing IL32β or a fLuc control. Respective genes were overexpressed at the time of IFNγ stimulation. Relative *Cm* infectivity is shown. **e.** An ELISA (n=3) to measure IL32 was performed on supernatant and lysate (in equivalent volume) from A549 and HeLa cells. IL32 knockout supernatant and lysate was used to subtract background signal. “N.D” = not detected. Data in **b** and **d** were analyzed with a Brown-Forsythe and Welch one-way ANOVA with Dunnett’s T3 multiple comparisons test. Any groups that do not share a letter are statistically different; p<0.05.

**Figure S2.**
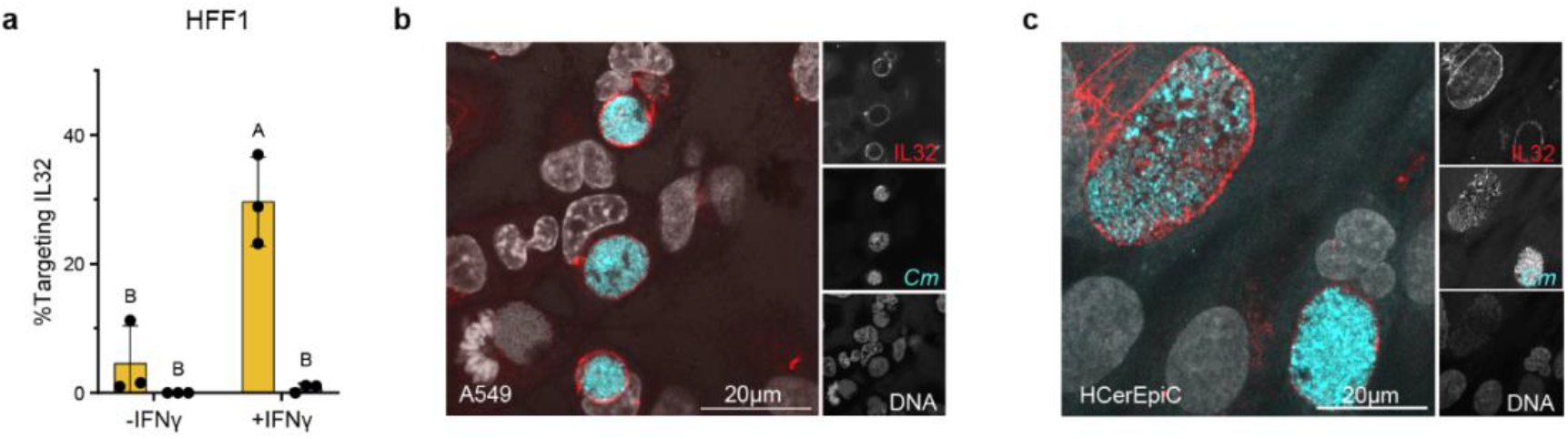
Intracellular IL32 targets *Cm* inclusions in epithelial cells and fibroblasts. **a.** Quantification of IL32 targeting to observed *Cm* or *Ct* inclusions in IFNγ-primed or unprimed HFF-1 cells. **b.** Representative image of IL32 targeting to *Cm* in A549 cells and **c.** in HCerEpiC primary cervical epithelial cells. Data in **a** were analyzed with a two-way ANOVA with Tukey’s multiple comparisons test.

**Figure S3.**
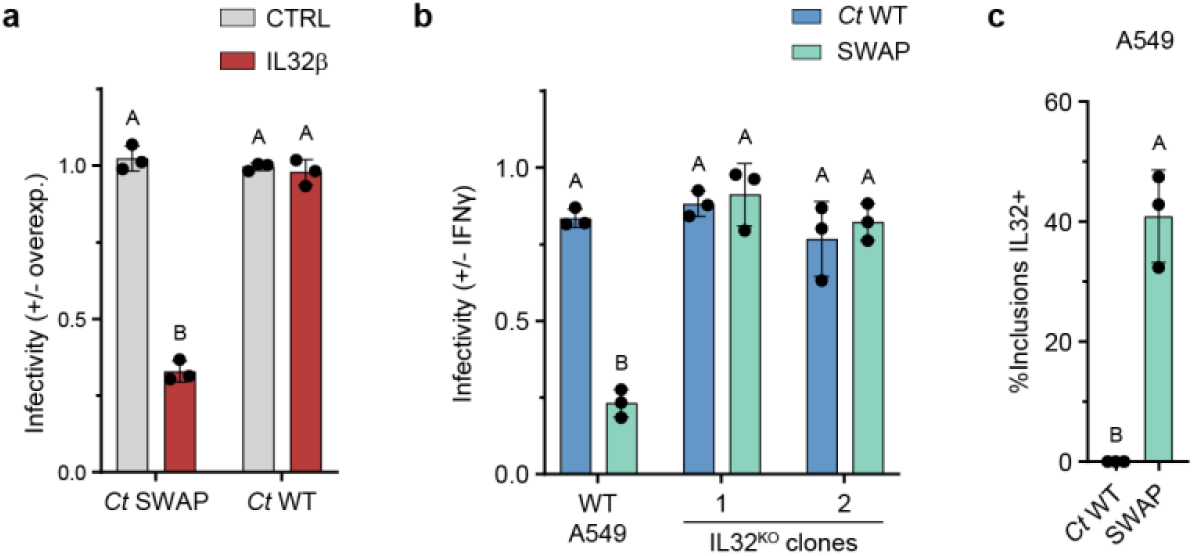
The *Ct* SWAP strain carrying the *Cm* IncS locus is susceptible to IL32. **a.** Relative infectivity of *Ct* WT and SWAP in A549 cells overexpressing IL32β or a fLuc control. **b.** Relative infectivity of *Ct* WT and SWAP in IFNγ-primed A549 cells (wild-type and IL32^KO^ clones 1 and 2). **c.** Quantification of IL32 targeting to observed *Ct* WT and SWAP inclusions in IFNγ-primed A549 cells. Data in **a** and **b** were analyzed with a two-way ANOVA with Tukey’s multiple comparisons test. Data in **c** were analyzed with an unpaired, two-tailed Welsh’s t-test.

**Figure S4.**
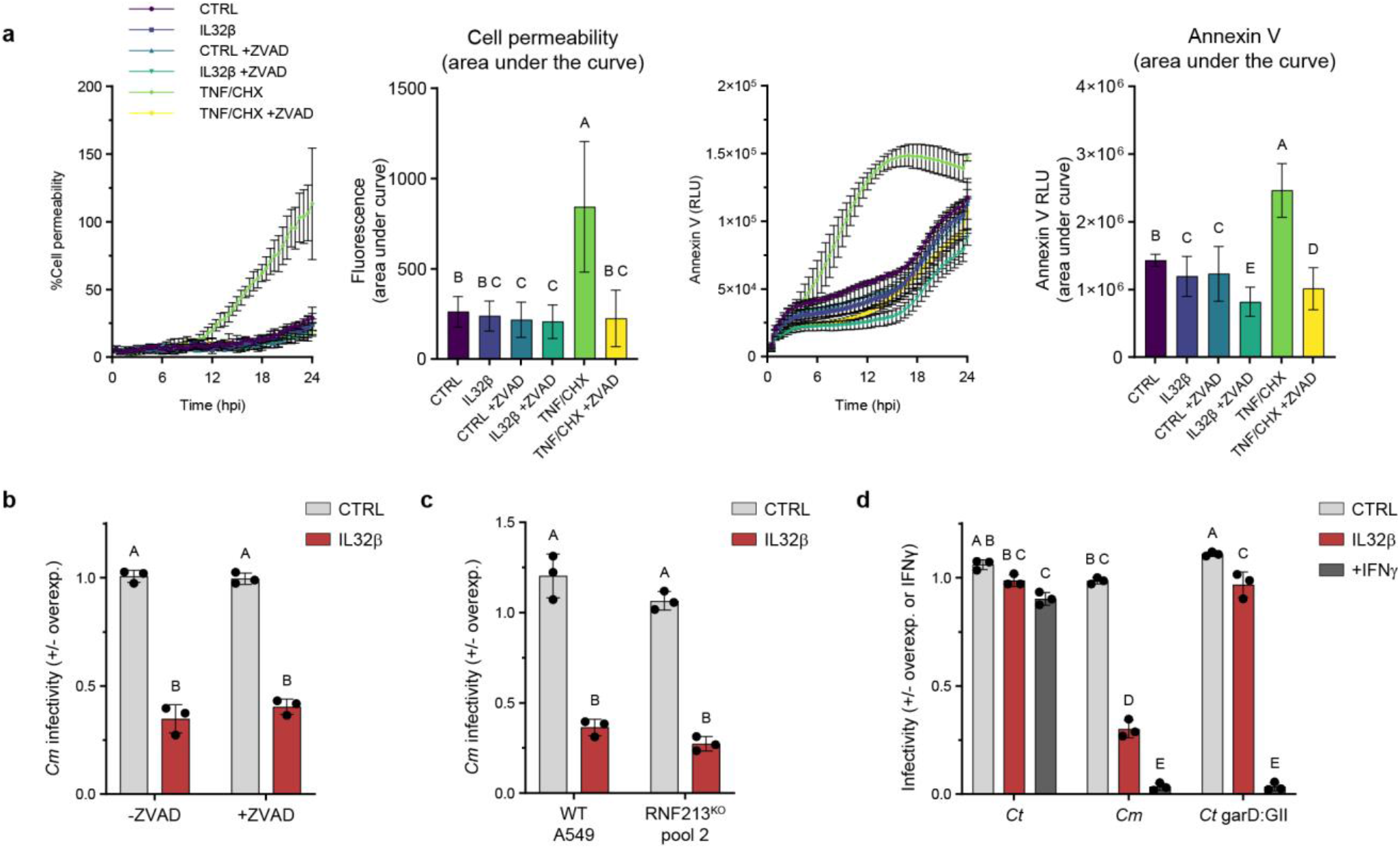
IL32 restricts *Cm* independently of cell death and the E3 ligase RNF213. **a.** Cell death over 24hpi during *Cm* infection. A549 cellss were pre-treated with 2µg/mL anhydrotetracycline to induce expression of IL32β or a fLuc control. After 24 hours, cells were infected with *Cm* at an MOI of 2 and treated with 20µM Z-VAD-FMK or left untreated. In parallel, untreated, uninfected A549s were treated with TNFα and cycloheximide, with or without Z-VAD-FMK. The RealTimeGlo kit was used to measure cell permeability (left, representing late-stage cell death) and annexin V exposure (right, representing early apoptosis). Both timecourse and “area under the curve” data are shown. **b.** A549 cellss overexpressing IL32β or a fLuc control were infected with *Cm* and treated with 0 or 20 µM Z-VAD-FMK. Relative infectivity is shown. **c.** Relative infectivity of *Cm* in wild-type or RNF213^KO^ pool 2 A549 cells overexpressing IL32β or a fLuc control. **d.** Infectivity of *Cm*, *Ct*, and a *Ct* garD mutant in A549 cells overexpressing IL32β, overexpressing a fLuc control, or treated with IFNγ. Data in **a** were analyzed with a Brown-Forsythe and Welch one-way ANOVA with Dunnett’s T3 multiple comparisons test. Data in **b-d** were analyzed with a two-way ANOVA with Tukey’s multiple comparisons test.

**Figure S5.**
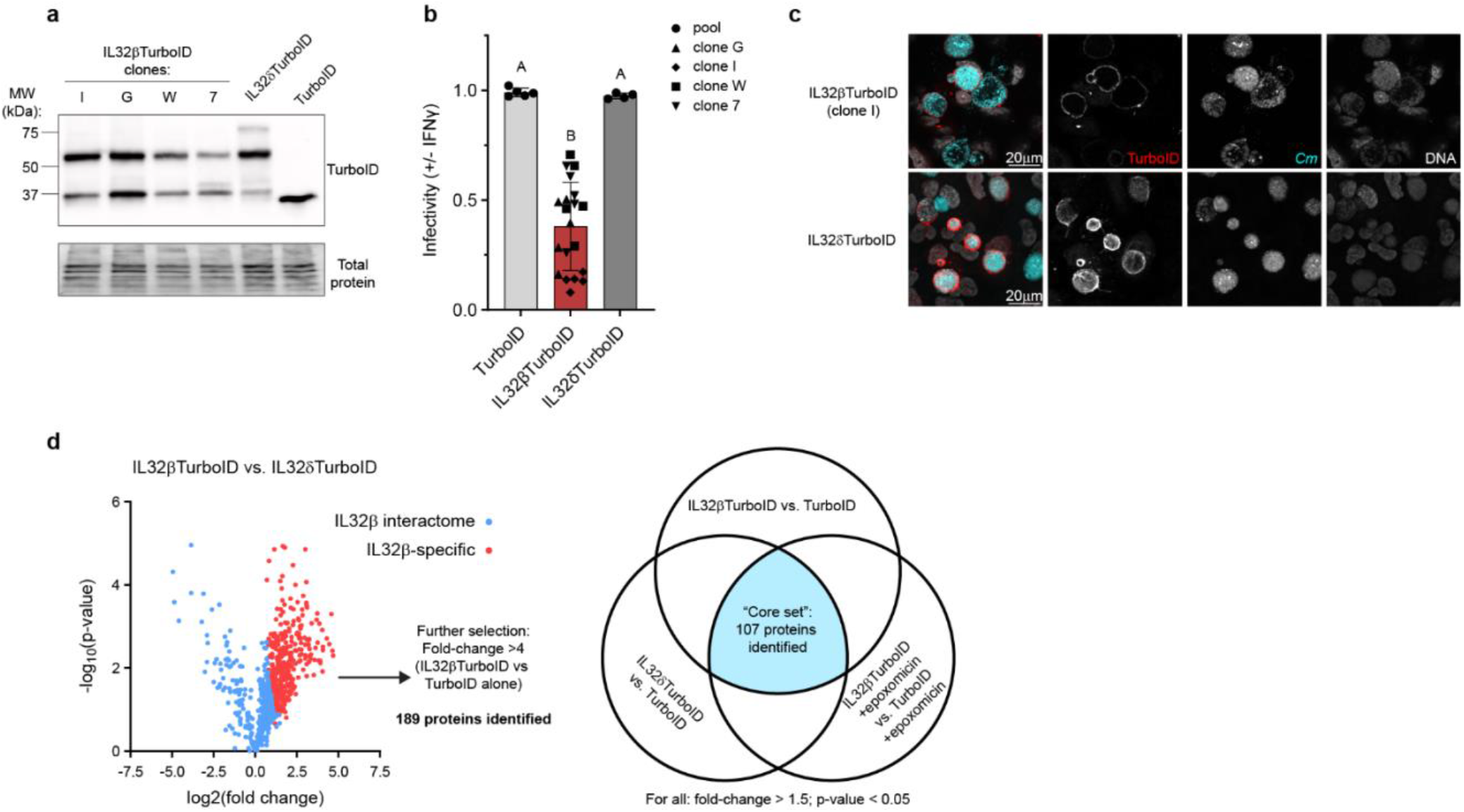
Validation and screen design with IL32-TurboID constructs. **a.** Western blot measuring TurboID and total protein from lysate of A549 cells overexpressing IL32βTurboID, IL32δTurboID, or TurboID alone. **b.** Relative infectivity of *Cm* in A549s overexpressing IL32βTurboID, IL32δTurboID, or TurboID alone. **c.** Representative images of IL32βTurboID and IL32δTurboID targeting *Cm* in A549 overexpression cells. **d.** Depiction of how protein “hits” from this TurboID screen were chosen for further analysis. Data in **b** were analyzed with a Brown-Forsythe and Welch one-way ANOVA with Dunnett’s T3 multiple comparisons test.

**Figure S6.**
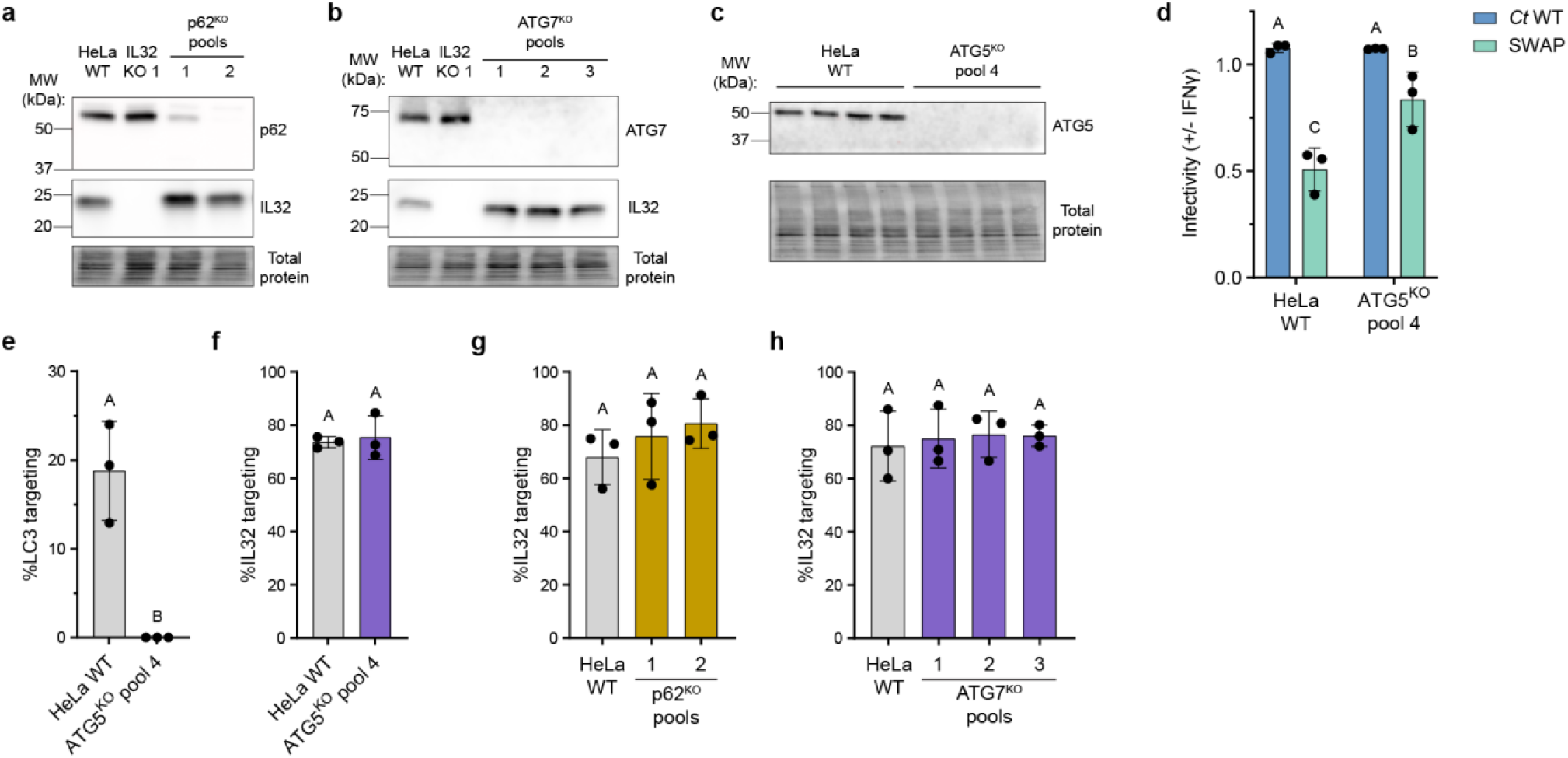
Validation and characterization of p62, ATG7, and ATG5 knockout cell lines. **a.** Western blot of p62, IL32, and total protein in lysates from IFNγ-primed p62^KO^ HeLa pools, with WT and IL32^KO^ controls. **b.** Western blot of ATG7, IL32, and total protein in lysates from IFNγ-primed ATG7^KO^ HeLa pools, with WT and IL32^KO^ controls. **c.** Western blot of ATG5 and total protein in lysates from WT and ATG5^KO^ pool 4 HeLa cells. **d.** Relative infectivity in IFNγ-treated HeLa cells (WT and Atg5^KO^ pool 4) infected with *Ct* WT or SWAP. **e.** Quantification of LC3 targeting and **f.** IL32 targeting to *Ct* SWAP in IFNγ-primed HeLa cells (WT and Atg5^KO^ pool 4). **g.** Quantification of IL32 targeting to *Ct* SWAP in IFNγ-primed HeLa cellss (WT and p62^KO^ pools). **h.** Quantification of IL32 targeting to *Ct* SWAP in IFNγ-primed HeLa cells (WT and ATG7^KO^ pools). Data in **d** were analyzed with a two-way ANOVA with Tukey’s multiple comparisons test. Data in **e** and **f** were analyzed with an unpaired, two-tailed Welsh’s t-test. Data in **g** and **h** were analyzed with a Brown-Forsythe and Welch one-way ANOVA with Dunnett’s T3 multiple comparisons test.

**Figure S7.**
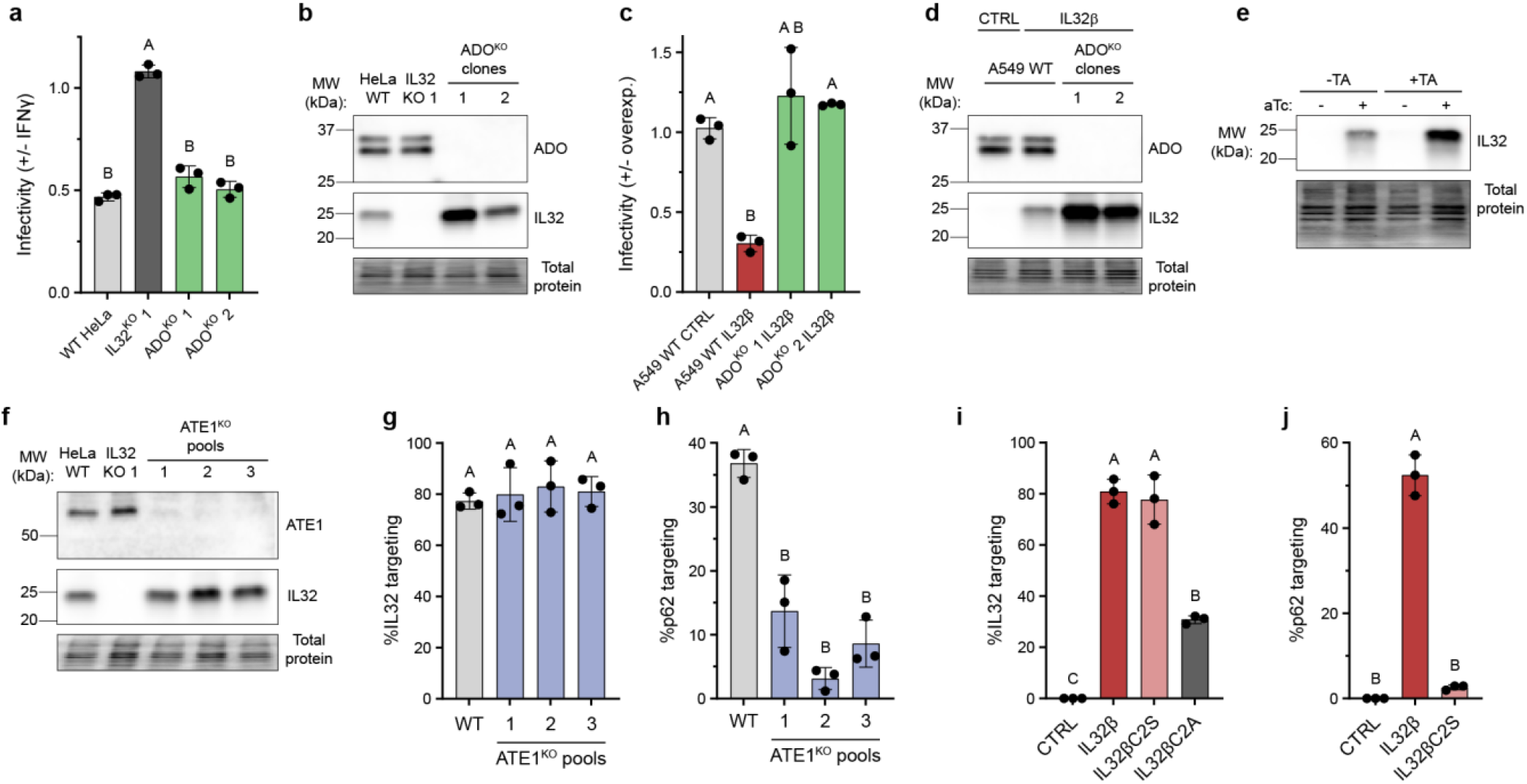
Validation of ADO^KO^ and ATE1^KO^ cells and their role in cell-autonomous defense against *Chlamydia*. **a.** Relative SWAP infectivity in IFNγ-primed ADO^KO^ HeLa clones, with WT and IL32^KO^ controls and **b.** Western blot of ADO, IL32, and total protein in lysates from IFNγ-primed ADO^KO^ HeLa clones with WT and IL32^KO^ controls. **c.** Relative SWAP infectivity in WT A549 cells or ADO^KO^ clones expressing IL32β or fLuc control. **d.** Western blot of ADO, IL32, and total protein of lysates from WT A549 cells or ADO^KO^ clones overexpressing IL32β or a fLuc control. **e.** Western blot of IL32 and total protein of lysates from WT A549 cells expressing IL32β, treated with 0 or 80µM tannic acid. **f.** Western blot of ATE1, IL32, and total protein in lysates from HeLa WT, IL32^KO^, ATE1^KO^ pools. **g.** Quantification of IL32 targeting and **h.** p62 targeting to *Ct* SWAP in IFNγ-primed WT or ATE1^KO^ pools HeLa cells. **i.** Quantification of IL32 targeting and **j.** p62 targeting to *Ct* SWAP in A549 IL32^KO^ clone 1 expressing IL32β, IL32βC2S mutant, IL32βC2A mutant, or a no vector control. Data in **g** and **h** correspond to Fig. 5d and data in **i** and **j** correspond to Fig. 5g. Data in **a**, **c**, **g**, and **h** were analyzed with a Brown-Forsythe and Welch one-way ANOVA with Dunnett’s T3 multiple comparisons test. Data in **i** and **j** were analyzed with a one-way ANOVA with Tukey’s multiple comparison’s test.

## MATERIALS AND METHODS

### Cell culture

A549 (ATCC #CCL-185), HeLa (ATCC #CCL-2), HEK-293T/17 (ATCC # CRL-11268) and Vero (ATCC #CCL-81) cells were cultured in Dulbeco’s Modified Eagle Medium (DMEM; Gibco) supplemented with 1% MEM Non-essential Amino Acids (NEAA; Gibco) and either 9% heat-inactivated fetal bovine serum (FBS; Omega Scientific) or 9% Cosmic Calf serum (Cytiva). Primary cervical epithelial cells (HCerEpiC; ScienCell, 7060) were kindly provided by Dr. Carolyn Coyne (Duke University) and cultured in EpiCM media (ScienCell) supplemented with EpiCGS (ScienCell) and 9% heat-inactivated FBS. E6/E7-immortalized oviduct epithelial cells were kindly provided by Dr. Catherine O’Connell (UNC, Chapel Hill) and cultured in Dulbecco’s Modified Eagle Medium/Ham’s Nutrient Mixture F-12 including GlutaMAX (DMEM/F12, Gibco) was further supplemented with 1% MEM Non-essential Amino Acids and 9% Cosmic Calf serum. All cells were grown at 37°C with 5% CO_2_ and were routinely tested for *Mycoplasma* contamination.

### Plasmids, cloning, and stable cell line production

pTRIP plasmids containing IL32 (isoform γ), firefly luciferase, and TNFα ORFs were kindly provided by Dr. Neal Alto (UT Southwestern). pcDNA 3.1 plasmids containing FLAG-tagged IL32γ and β were generated by VectorBuilder and kindly provided by Dr. Carolyn Coyne (Duke University). These ORFs (IL32γ, firefly luciferase, TNFα, IL32γ-FLAG, and IL32β-FLAG) were PCR amplified using attB-flanked primers and subsequently gateway cloned into the pDONR221 backbone (Invitrogen) using BP Clonase II (Thermo) following manufacturer’s instructions. Once in pDONR221, the untagged IL32γ construct was mutated using the Q5 site-directed mutagenesis (SDM) kit (NEB) to produce pDONR221-IL32β. This IL32β construct was further mutated using Q5 SDM to produce Cys2Ala and Cys2Ser mutants. The IL32β-FLAG construct was mutated using Q5 SDM to produce FLAG-tagged IL32δ, IL32 isoform 5, and IL32 isoform 6. IL32 isoform 5 was then further mutated using Q5 SDM to produce FLAG-tagged IL32 isoform 7. To produce IL32β-TurboID fusion protein, we used the InFusion HD Cloning Kit (Takara) following manufacturer’s instructions. Briefly, we PCR amplified pDONR221-IL32β, amplified TurboID template from a plasmid kindly gifted by Dr. Scott Soderling (Duke University), gel purified these PCR products (Takara), and combined PCR products using the InFusion cloning kit. Once pDONR221-IL32βTurboID was produced, we subsequently produced pDONR221-IL32δTurboID and pDONR221-TurboID using Q5 SDM.

All of the above constructs in pDONR221 were subsequently cloned into pInducer20^44^ (a gift from from Stephen Elledge; addgene #44012) using LR Clonase II (Thermo) and following manufacturer’s instructions. pInducer20 plasmids were packaged into lentivirus using psPAX2 (a gift from Didier Trono; addgene #12260) and VSVG for packaging. HEK-293T cells were transfected with these vectors using TransIT-293 transfection reagent (Mirus). Virus-containing media was collected at 48- and 72-hours post transfection, pooled, filtered through 0.45 μm nylon filter (Corning), aliquoted, and stored at -80°C. A549 cells were transduced through spinning at 1000xg for 45 minutes at room temperature with freshly thawed virus. To establish stable cell lines, cells were treated with 2mg/mL geneticin (Gibco) starting 48 hours after transduction, which was maintained until all untransduced control cells were dead.

The p62 constructs used in Fig. 5J were gifts from Benjamin Wolozin^45^. “CTRL” is pHR-SFFV-FLAG-TurboID, addgene #223713. “WT” is pHR-SFFV-FLAG-TurboID-SQSTM1, addgene #223714. “PB1” is pHR-SFFV-FLAG-TurboID-dPB1-SQSTM1, addgene #223716. “LIR” is pHR-SFFV-FLAG-TurboID-mutLIR-SQSTM1, addgene #223718. “UBA” is pHR-SFFV-FLAG-TurboID-dUBA-SQSTM1, addgene #223717. To generate a “ZZ” mutant, Q5 site-directed mutagenesis was performed on WT plasmid #223714 to remove amino acids 123-173 of p62. As with the pInducer20 constructs, lentiviral vectors were produced in HEK-293T cells and similarly transduced into HeLa cells. All newly generated plasmids in this study have been confirmed with Sanger sequencing analysis (Plasmidsaurus or GeneWiz).

### Chlamydia strains

All *C. trachomatis* strains used were in a serovar LGV-L2/434/Bu background. For visualization, *C. trachomatis* strains were previously transformed with pGFP::SW2^46^ or p2TK2-mCherry^47^. *C. muridarum* MoPn strains with or without a pGFP::CM plasmid^8,48^ were used as well. For co-infection studies, a previously described *incA* nonsense mutant strain of *C. trachomatis* was used^15,49^. For some studies demonstrating RNF213 independence, a previously described *Ct* garD::GII strain was used^9,50^.

*C. trachomatis-C. muridarum* chimera strains RC13, RC3834, RC4243, RC108, RC1696, RC1203, RC27, RC1420, RC826, RC1323, RC1219, RC2043, RC2457, RC26, RC6882, RC3772, RC2180, RC2220, RC302, and RC6389 were previously developed through a lateral gene transfer approach^16,17^. *C. trachomatis* strains used for IncS validation studies include the previously established IncS “SWAP” strain^19^ and the IncS “cKO” strain^18^.

### *Chlamydia* propagation and infection

*Chlamydia* infections were performed as previously described^9^. Briefly, fresh aliquots of *Chlamydia* seed preps were thawed at room temperature, diluted in cold cell culture medium, and added to cell monolayers. Cells were then spun at 3000 rpm at 4°C for 30 minutes in a Sorvall ST 40R centrifuge and were then returned to a 37°C incubator. “Time of infection” was defined as the halfway point (15 minutes) into the 30-minute centrifugation step. All infections were performed at a multiplicity of infection (MOI) of 2 except for co-infection assays (MOI 3 for *Cm*-GFP, MOI 5 for *Ct* incA mutant) and the proteomics screen (MOI of 8 for *Cm*-GFP).

*Chlamydia* spp. were grown in confluent Vero cells for 26 hours (*Cm*), 48 hours (*Ct* and *Cm-Ct* chimeras), or 40 hours (*Ct* IncS SWAP) before washing samples once in UltraPure Distilled Water (diH_2_O; Invitrogen) and lysing in diH_2_O for 20 minutes with scraping and pipetting. 5X SPG was then added to reach a final concentration of 1X (final concentration: 0.5 g/L KH_2_PO_4_, 1.2 g/L Na_2_PO_4_, 0.72 g/L L-glutamic acid, 75 g/L sucrose, pH 7.5), and *Chlamydia* was subsequently aliquoted and stored at -80°C until use. For each seed prep, a bacterial titer was determined by thawing and serially diluting one aliquot, infecting confluent Vero cells, and manually counting inclusions in a moderately infected well.

### *Chlamydia* infectivity assays

Bacterial burden assays were performed as previously described^9^. Cells were seeded at 0.8-1.2×10^4^ cells per well in a clear or black, clear-bottom 96-well flat-bottom plates (Corning). The next day, cells were primed with 0-100U/mL IFNγ (Millipore) 16 hours before infection, or with 0-2µg/mL anhydrotetracycline (Takara) 24h before infection to induce overexpression of pInducer20^44^ constructs. Cells were then infected with an MOI of 2 (2-4×10^4^ IFU/well). All priming and infections were performed in appropriate cell culture media supplemented with 100µg/mL L-Trp unless otherwise stated. 24 hours after infection, cells were fixed in cold 4% paraformaldehyde (PFA) in PBS (Gibco) for 20 minutes.

Fluorescent *Chlamydia* strains were stained with 1:1000 Hoechst 33258 (Invitrogen) in PBS for at least 10 minutes. Non-fluorescent *Chlamydia* strains were permeabilized in ice-cold methanol for 1 minute, blocked in blocking buffer (5% bovine serum albumin, 2% glycine in PBS) for 30 minutes, and stained in primary 1:1000 anti-MOMP (Medix Biochemica) followed by secondary 1:1000 anti-goat AlexaFluor488 and Hoechst 33258.

Both nuclei (DAPI channel) and inclusions (GFP or mCherry channel) were imaged and counted by the CellInsight CX5 HCS Platform (Thermo). We first determined the number of Chlamydial inclusions per host nuclei for each well using a custom thresholding algorithm, then averaged at least two technical replicate wells per group. “Relative infectivity” was calculated by taking the average inclusions per nuclei of a treated group (e.g. +IFNγ) and dividing it by the average inclusions per nuclei of an untreated control (e.g. -IFNγ).

### Immunofluorescence and targeting assays

Cells were seeded on 12 mm #1.5 glass coverslips (Fisher) at 1.5-2.0×10^5^ cells/well. The next day, cells were infected with 2 MOI (3.0-4.0×10^5^ IFU/well) of *Chlamydia*. Three hours post infection, cells were primed with 0 or 100U/mL IFNγ, or 0 or 0.2µg/mL anhydrotetracycline (Takara). All priming and infection steps were performed in appropriate cell culture media supplemented with 100µg/mL L-Trp. 20 hours after infection, cells were washed twice in sterile PBS (Gibco), fixed in cold 4% paraformaldehyde (PFA; Sigma) in PBS for 20 minutes, and washed twice in PBS.

Cells were permeabilized through one of three methods: 1) ∼1-minute treatment in ice-cold 100% methanol, followed by two PBS washes, 2) inclusion of 0.05% saponin (Amresco) throughout staining, or 3) a 15-minute treatment in 0.1% Triton X-100 (ProPure) in PBS, followed by the inclusion of 0.05% Triton X-100 in all wash steps. In all cases, coverslips were blocked for 30 minutes in blocking buffer (5% bovine serum albumin (VWR), 2.2% L-Glycine (VWR), +/- 0.05% saponin in PBS). Coverslips were then stained for 1 hour in primary antibody diluted in blocking buffer, then washed three times in wash buffer (PBS +/- 0.05% saponin, +/- 0.05% Triton X-100). Coverslips were then stained for 1 hour in diluted secondary antibody in blocking buffer, then washed an additional three times in wash buffer. Finally, coverslips were plated on microscopy slides in Mowoil mounting media (Sigma) with 0.01% p-phenylenediamine (Sigma) and left to dry overnight.

Coverslips were imaged on a Zeiss Axio Observer Z1 with a 63x oil objective. Five to seven images were taken per coverslip (typically 100-200 *Chlamydia* inclusions per group) unless otherwise stated and groups were either blinded manually or with the FIJI “File Name Encrypter” Plug-In Tool. For each image, the FIJI “Cell Counter” tool was used to count the total number of *Chlamydia* inclusions and the number of inclusions targeted by each factor. For all proteins “targeting” was defined as a clear ring that is visibly brighter than background signal. For all proteins except for LC3, this ring must surround or cover at least 50% of the inclusion. For LC3, this ring must surround at least 20% of the inclusion. For some experiments, additional representative images were taken with a Zeiss Axio Observer Z1 with Zeiss Apotome 3, or with a Zeiss 880 Airyscan inverted confocal microscope.

### *Microsporidia* strains and propagation

*Encephalitozoon cuniculi* strains ECI and ECII^51^ were a gift from Dr. Edward Miao (Duke University). Strains were propagated in Vero cells for 7-12 days before harvesting spores, with media replaced every 2-3 days. To harvest spores, 10 cm dishes containing infected Vero cells were washed once with sterile water, then incubated in 10 mL water until Vero cells were lysed (5-10 minutes). Dishes were scraped with cell lifters, then pipetted up and down 5-10 times to further lyse the Vero cells. Lysate was filtered through a 5µm syringe filter (PALL). Spores were pelleted by centrifugation at 2,000xg for 10 minutes, washed once in sterile water and centrifuged again at 2,000xg for 10 minutes. Spores were then resuspended in 1mL PBS and stored at 4°C until infection. Number of spores per mL was quantified using a hemocytometer.

### *Microsporidia* infectivity and immunofluorescence assays

Infectivity and immunofluorescence assays for *Microsporidia* strains ECI and ECII were almost identical to those for *Chlamydia*. For infectivity assays, plating and priming were identical. Cells were infected with an MOI of 20 (4×10^5^ IFU/well) for ECI and ECII, and 0 or 2µg/mL anhydrotetracycline was maintained throughout infection to preserve expression of pInducer constructs. Infections were performed by spinning cells at 800xg at room temperature for 10 minutes before returning cells to the 37°C, 5% CO_2_ incubator. As with *Chlamydia* infections, all priming and infection steps were performed in cell culture media supplemented with 100µg/mL L-Trp. At 50hpi, cells were washed twice with sterile PBS and fixed for 20 minutes with cold 4% PFA in PBS for 20 minutes. Cells were then washed twice in PBS, permeabilized for 15 minutes in 1% Triton X-100 in PBS, and then stained in 1:1000 Calcofluor white (Biotium) to stain Microsporidial spores and 1:10,000 SYTOX green (Thermo) to stain nuclei for 1 hour. Cells were then washed again in PBS. Both nuclei (green channel) and microsporidia (blue channel) were imaged and counted by the CellInsight CX5 HCS Platform (Thermo). “Relative infectivity” was calculated the same as for *Chlamydia*.

For immunofluorescence assays, cells were plated on coverslips identically to *Chlamydia* protocols. One day after plating, cells were infected with an MOI of 10 (4×10^6^ IFU/well) of ECI or ECII. At 26hpi, cells were primed with 0 or 100U/mL IFNγ. At 50hpi, cells were washed twice in sterile PBS, fixed in 4% PFA for 20 minutes, and washed twice more in PBS. Coverslips were stained following the above Triton X-100 permeabilization protocol, and using both Calcofluor white and SYTOX green at the above concentrations to stain ECI/ECII and nuclei, respectively. For the representative image in Fig. 2f, we used a custom rat antibody against *E. cuniculi* NTT2, which was a gift from Dr. Robert Hirt (Newcastle University)^52^.

### Western blot

Cells were seeded at 2.0×10^5^ cells/well of a clear 24-well plate (Corning). The next day, cells were primed with 0-100U/mL IFNγ (Millipore) 16 hours before lysis, or with ≤2µg/mL anhydrotetracycline (Takara) 24h before lysis to induce overexpression of pInducer20^44^ constructs. Cells were washed twice in sterile PBS (Gibco) and then lysed for 30 minutes at 4°C in Pierce RIPA buffer (Thermo) containing 1:100 DNase I (NEB) and protease inhibitor cocktail (Sigma). Lysate was then collected, clarified by spinning at 10,000xg for 10 minutes, added to 4x Laemmli buffer (BioRad) +10% β-mercaptoethanol (VWR) to a final concentration of 1x, and heated at 70°C for 10 minutes or 95°C for 5 minutes.

Samples were run on 4-20% Stain-Free gels (BioRad) and gels were activated with UV for 5 minutes to visualize total protein. Protein was transferred onto methanol-blocked Immobilon FL PVDF membranes (Millipore) using a Trans-Blot Turbo Transfer System (BioRad). Unless otherwise stated, membranes were stained overnight in primary antibody in blocking buffer (5% nonfat dry milk (BioRad) in Tris-buffered saline including 0.1% Tween-20 (TBST)). Membranes were then washed three times for five minutes in TBST, stained for one hour in 1:5000 secondary antibody in blocking buffer, and washed three times for seven minutes in TBST. Depending on signal intensity, membranes were imaged in one of three ECL solutions on an Azure 500 Western Blot imager (Azure Biosystems): Clarity ECL Substrate (BioRad), Amersham ECL Prime Western Blotting Detection Reagent (Cytiva), or SuperSignal West Femto ECL substrate (Thermo Fisher). If necessary, membranes were stripped for 30 minutes at room temperature with Restore Western Stripping Buffer (Thermo) followed by five washes in TBST and an additional primary antibody incubation.

### Supernatant transfer, ELISA, and cell death assay

For the supernatant transfer and ELISA experiments, cells were plated and primed in 96 well plates following an identical protocol to other *Chlamydia* infectivity assays. 16 hours after IFNγ priming, or 24 hours after pInducer-mediated overexpression, supernatant was removed from cells for supernatant transfer or ELISA. For the supernatant transfer experiment (Fig. 1j), this supernatant was applied to fresh, wild-type A549 cells plated the previous day. These cells were incubated for 24 hours at 37°C in transferred supernatant, and then were infected, fixed, stained, and imaged following the *Chlamydia* infectivity assay protocol. For the ELISA, supernatant was collected and cells were lysed for 5 minutes in an equivalent volume of cell culture medium including 1% Triton X-100 (ProPure). IL32 was measured in supernatant and lysate using the IL32 DuoSet ELISA kit (R&D systems) following manufacturer’s instructions and using a CLARIOstar Plus plate reader. To eliminate non-specific background signal, supernatant and lysate absorbances from A549 and HeLa cells was subtracted by corresponding supernatant and lysate absorbances from IL32 knockout A549 and HeLa controls.

Cell death assays were performed with the Real-Time Glo Annexin V Apoptosis and Necrosis kit (Promega) following manufacturer’s instructions. A549 cells were plated, primed, and infected following an identical protocol to other *Chlamydia* infectivity assays, except white, clear bottom 96-well plates (Corning) were used and all steps were performed in phenol red-free DMEM containing 9% heat inactivated FBS and non-essential amino acids. At the time of infection, Z-VAD-FMK (Invivogen, final concentration 20µM) was added to some wells to block caspase-mediated cell death, a combination of TNFα (R&D Systems, final concentration 50ng/mL) and cycloheximide (Fluka, final concentration 10µg/mL) was added to some wells to induce apoptosis, and 1% Triton X-100 was added to some wells to induce total cell lysis. At the time of infection, all wells were treated with 1X Real-Time Glo detection reagents, and the 96 well plate was moved to a CLARIOstar Plus plate reader with incubation at 37°C with 5% CO_2_. Fluorescence and luminescence were measured every 30 minutes for 24 hours.

### ISG screen

Lentiviral particles were kindly provided by Neal Alto (UT Southwestern) and produced as previously described^10,53^. Candidate type II interferon-stimulated genes (ISGs) were compiled from published transcriptomic datasets of various human tissues and cell lines stimulated with IFNγ. Genes exhibiting greater than 2-fold increases in expression (with statistical significance in each study’s analysis) in two or more datasets were cloned from Ultimate ORF Collection (Thermo) by Dr. John Schoggins (UT Southwestern), yielding 470 genes for library construction. ORFs, along with a bicistronic tagRFP containing its own internal ribosome entry site, were cloned into the pTRIP plasmid vector and driven by a CMV promoter for overexpression. Firefly luciferase (fLUC) was used as a negative control. The type II ISG lentiviral library was prepared as previously described^10,53^. Briefly, 293T cells were transfected with 1μg pTRIP plasmid, 0.8μg pGAG-pol and 0.2μg VSVg. The next day, media was changed to fresh “pseudoparticle” DMEM (DMEM supplemented with 3% heat-inactivated FBS, 1% NEAA and 20mM HEPES (Gibco)). Virus was harvested and pooled after an additional 24 and 48 hours and supernatant was filtered through 0.45µM filters (Corning), supplemented with 4 μg/mL polybrene and stored in 96-well plates.

A549 cells were plated as described above. The next day, library plates were thawed on ice and cells were transduced with 25 μL lentivirus in 150 μL “pseudoparticle” DMEM containing 4 μg/mL polybrene and 100μg/mL L-Trp. Plates were spun at 37°C, 1000 g for 1 hour. 24 hours after transductions, plates were infected with *Cm* at an MOI of 2 and assayed for infectivity as described above. Screen results were assessed statistically by Z-score, and “hits” were classified as ISGs displaying a Z-score less than -2 (p < 0.05) in two independent screening attempts. pTRIP plasmids of hits were used to make independent batches of lentivirus for validation and sequencing confirmation.

### Proteomics screen

A549 cells were plated at 8.0×10^5^ cells/well in 6 well plates (Corning). These cells were either pools containing pInducer20-TurboID or pInducer20-IL32δTurboID; or four clones containing pInducer20-IL32βTurboID. The next day, to normalize expression of each construct, IL32βTurboID cells were treated with 2µg/mL anhydrotetracycline (aTc), IL32δTurboID cells were treated with 0.75µg/mL aTc, and TurboID cells were treated with 0.1µg/mL aTc. 24 hours later, all cells were infected with an MOI of 8 (9.6 million IFU/well) of *Cm*. To some wells, a final concentration of 100nM epoxomicin (MedChemExpress) was added to preserve short-lived interactors. 4 hours post-infection (hpi), biotin (Sigma) was added to all groups at a final concentration of 50µM. At 9hpi, cells were washed five times in sterile PBS, and then lysed for 30 minutes at 4°C in RIPA buffer containing 1:100 DNase I and protease inhibitor cocktail. Lysate was then collected and clarified by spinning at 10,000xg for 10 minutes. Protein quantity for each sample was measured using the Pierce BCA Protein Assay Kit (Thermo) following manufacturer’s instructions.

To purify biotinylated proteins from lysate, MagStrep Strep-Tactin beads (IBA) were used following the manufacturer’s protocol. Briefly, 1mg of protein per sample was added to 60µL beads and left rocking for 16 hours at 4°C. For IL32βTurboID cells, 250µg protein per clone was used, to a total of 1mg. Beads were washed four times in wash buffer (0.1M Tris-Cl, 0.15M NaCl, 1mM EDTA, 0.4% SDS) and were separated from washes using a DynaMag-2 magnetic separation rack (Invitrogen). Protein was eluted in 2x laemmli buffer (65.8mM Tris-HCl, 26.3% (w/v) glycerol, 2.1% SDS) + 20mM DTT (Roche) through vigorous vortexing and heating to 95°C for ten minutes. Beads were removed twice for each sample with a magnetic separator. Collected protein was stored at -80°C and three repeats per group were given to the Duke Proteomics Core Facility for LC-MS/MS Analysis.

Quantitative LC/MS/MS was performed using an EvoSep One UPLC coupled to a Thermo Orbitrap Astral high resolution accurate mass tandem mass spectrometer (Thermo). Data collection on the Orbitrap Astral mass spectrometer was performed in a data-independent acquisition (DIA) mode of acquisition with a r=240,000 (@ m/z 200) full MS scan from m/z 380-980 in the OT with a target AGC value of 4e5 ions. Fixed DIA windows of 4 m/z from m/z 380 to 980 DIA MS/MS scans were acquired with a target AGC value of 5e4 and max fill time of 6 ms. Data were then imported into Spectronaut (Biognosis) and individual LCMS data files were aligned based on the accurate mass and retention time of detected precursor and fragment ions. Relative peptide abundance was measured based on MS2 fragment ions of selected ion chromatograms of the aligned features across all runs. The MS/MS data was searched against SwissProt *Homo sapiens* and *Chlamydia muridarum* databases, a common contaminant/spiked protein database (bovine albumin, bovine casein, yeast ADH, etc.), and an equal number of reversed-sequence “decoys” for false discovery rate determination. A library free Direct DIA+ approach within Spectonaut was used to perform these database searches.

To identify a first pass of proteins enriched in IL32βTurboID compared to TurboID alone, a two-tailed heteroscedastic T-test was performed on log_2_-transformed protein expression data. Using a fold-change cutoff of >1.5, an unadjusted p-value cutoff of <0.05, and a precursor cutoff of >1, we identified 844 human and *Cm* proteins enriched in the IL32βTurboID group compared to TurboID alone, hereafter referred to as the “IL32β interactome”. We identified two subsets of this interactome for further analysis. The first are “IL32β-specific”, or proteins that are enriched in IL32βTurboID compared to the IL32δTurboID control. Using MaxQuant Perseus software, we performed an additional T-test comparing the IL32βTurboID group to the IL32δTurboID group using a permutation-based FDR (fold-change>1.5, p-value<0.05 on log_2_transformed data). We further filtered the statistical hits of this analysis by removing *Cm* proteins, removing skin epithelial components KRT28 and SPRR1A, and removing proteins with a fold change <4 for the initial IL32βTurboID vs. TurboID comparison. This produced 189 “IL32β-specific” proteins. Our second subset of interest was the “core set” proteins, or proteins that are enriched in three settings: IL32βTurboID vs. TurboID alone, IL32δTurboID vs. TurboID alone, and IL32βTurboID + epoxomicin vs. TurboID alone + epoxomicin. For each comparison, we performed the same analysis as above, a heteroscedastic T-test on log_2_transformed protein expression data using a fold-change cutoff of >1.5 and an unadjusted p-value cutoff of <0.05. Only 107 human proteins, not including IL32 itself, were found to be significant in all three conditions. By combining the “β-enriched” and “core set” proteins, and removing duplicates, we identified 289 hits of our proteomic screen for further analysis.

### CRISPR-Cas9 genomic screen

Triple guide RNAs for each gene of interest were purchased from EditCo in a 384 well plate format. Additional control gRNAs were also purchased, namely positive controls IL32 and STAT1, and negative controls AAVS1 and scrambled control. A549 cells containing pInducer20-IL32β and wild-type HeLa cells were transduced with lentivirus containing the lentiCas9-Blast plasmid^54^ (addgene 52962) and were selected with blasticidin S (Invitrogen; 2.5µg/mL for HeLas, 7.5µg/mL for A549s) for five days until untransduced control cells were dead. In 96 well plates, an Echo 550 Acoustic Liquid Handler was used to dispense 2pmol per well of each guide RNA. On top of this RNA, 50µL of OptiMEM (Gibco) containing 0.2µL/well RNAiMAX transfection reagent (Invitrogen) was added. Finally, an additional 50µL of cell culture media containing 5×10^3^ cells/well was added, containing either HeLa-Cas9 cells or pInducer20-IL32β A549-Cas9 cells.

Cells were allowed to grow for three days to allow for Cas9-mediated knockout, and then HeLa cells were primed with 0 or 100U/mL IFNγ, and pInducer A549s were primed with 0 or 2µg/mL anhydrotetracycline (Takara). The following day, cells were infected with 2 MOI (4×10^5^ IFU/well) *Cm*-GFP, and 24 hpi cells were fixed in cold 4% paraformaldehyde (PFA) in PBS for 20 minutes. Subsequent Hoechst staining, high content imaging, and quantification were the same as described above. Figure 4d displays the average relative infectivity of three repeats in HeLa Cas9 cells and the average of two repeats in A549 pInducer20-IL32β Cas9 cells for each gRNA.

### Software and statistics

Graphs were generated and statistics were analyzed using GraphPad Prism 11. Unless otherwise stated, each experiment has an “n” of 3 or greater, where n is the number of independent experiments performed. Unless otherwise stated, significant results were defined as having a p-value less than 0.05 by an unpaired, two-tailed Welsh’s t-test, a Brown-Forsythe and Welch one-way ANOVA with Dunnett’s T3 multiple comparisons test, or a two-way ANOVA with Tukey’s multiple comparisons test.

Microscopy images were processed with FIJI ImageJ^55^. Diagrams in Figures 1c, 2i, 4a, 4c, and 5j were produced with BioRender. Figures were prepared using Adobe Illustrator.

### CRISPR-mediated knockout

A549 knockout pools for IL32 and RNF213 were generated by the Duke Functional Genomics Core using CRISPR/Cas9 technology as previously described^9^. Guide RNAs (gRNAs) targeting early exonic regions of each gene were designed using CHOPCHOP^56^ and Cas-OFFinder^57^ and cloned into PX459 V2^58^ (Addgene #62988). Cells were transfected with individual or paired sgRNAs using Lipofectamine 3000 (Invitrogen) or TransIT-LT1 (Mirus Bio) following manufacturer’s instructions. 24 hours after transfection, cells were selected with 2 μg/mL puromycin (Sigma) for three days, followed by growth as CRISPR-edited pools.

HeLa knockout pools for ATE1, ATG5, ATG7, SQSTM1 and IL32 were generated by the Duke Functional Genomics Core Facility. sgRNAs were designed using CHOPCHOP^56^ and ordered as modified synthetic sgRNAs from Synthego. 5 x 10^4^ HeLa cells were electroporated with 6 pmol TrueCut Cas9 protein v2 (Invitrogen) complexed with 18 pmol sgRNA using the Neon system (ThermoFisher Scientific) with the following settings: 1300 V, 20 ms, 2 pulses. Cells were recovered and expanded, followed by PCR sequencing and analysis using Inference of CRISPR Edits (ICE) to determine KO efficiencies for each individual sgRNA^59^.

For knockout cells in ADO (A549 and HeLa) and for HeLa p62 knockout clone 1, triple gRNA sets were designed and provided by EditCo, and were transfected into Cas9-expressing cells following the same protocol as the CRISPR genomic screen, except multiplying all quantities by five and transfecting in a 24 well plate (Corning). Knockout pools were subsequently expanded.

To generate knockout clones lacking ADO, IL32, p62, and RNF213; knockout pools were diluted to single cell concentrations, expanded, and genotyped by Western blotting. For all knockout cells, at least three confirmatory western blots were performed.

All guide RNAs used in this study are attached below:

ADO (in Cas9 background):

A549 clones 1/2; HeLa clones 1/2

- CGGCGCGGAGCUGGGUCAGG
- AAGCCUGGCGGGCGAUCCGU
- CGACGCGGCUUCUGGCCCGG

ATE1:

HeLa pool 1

- CCACAATCTGGCCCATTAGG

HeLa pool 2

- CTTCAGTCACAAGATTTCGT

HeLa pool 3

- TGAGTAACTCTTACCTCCAA

ATG5:

HeLa pool 4

- CTACCTGATATATTCTAAAG

ATG7:

HeLa pool 1

- CTTGAAAGACTCGAGTGTGT

HeLa pool 2

- GCTGCCAGCTCGCTTAACAT

HeLa pool 3

- CTTCCAAGGTCAAAGGACGA

IL32:

A549 IL32KO clone 1

- GACAGTGGCGGCTTATTATG
- GTGAGTATGACACACCCATC

A549 IL32KO clone 2

- AGGGAGGAGCATTACCATTC
- GCTTCTTCATGTCATCAGAG

HeLa IL32KO clones 1 and 3

- GTCCTACGGAGCCCCACGGG

HeLa IL32KO clone 2

- GCTTCTTCATGTCATCAGAG p62:

HeLa pool 1

- CGACTTGTGTAGCGTCTGCG

HeLa pool 2

- AGCCATCGCAGATCACATTG

HeLa clone 1:

- UAUGGCGUCGCUCACCGUGA
- CUGCAGCCCCGAGCCUGAGG
- CUGCGAGCGGCUGCUGAGCC

RNF213:

A549 pool 1 / clone 1:

- GTGGACCGATTTGCAGTACA

A549 pool 2 / clone 2:

- CACGTGGTACCATTGCCGGA

A549 pool 3

- CGTCTTCATCGGCTACCACT

RNF213/IL32 dKO A549:

Clone 1 (in IL32KO clone 1 background):

- GTGGACCGATTTGCAGTACA

Clone 2 (in IL32KO clone 2 background):

- CACGTGGTACCATTGCCGGA

### Data storage

Proteomics data and the numerical values of all other quantified data depicted in data panels in this manuscript will be made openly available in the Digital Repositories at Duke at the time at which the final version of the manuscript is published. Digital object identifier will be provided.

## REAGENT LIST

| Reagent or resource | Source | Catalog Number | Notes |
| --- | --- | --- | --- |
| **Primary antibodies** |  |  |  |
| Anti-ADO | Abcam | ab134102 | WB 1:2000 |
| Anti-ATE1 | Atlas Antibodies | HPA038444 | WB 1:2000 |
| Anti-ATG5 | Cell Signaling Technology | 2630S | WB 1:1000 |
| Anti-ATG7 | Cell Signaling Technology | 2631T | WB 1:2000 |
| Anti-Chlamydial MOMP | Medix Biochemica | V1621_1ml | IF 1:1000 (methanol) |
| Anti-EcNTT2 | Dr. Robert Hirt (Newcastle University) | N/A | IF 1:200 (triton) |
| Anti-FLAG (clone M2) | Sigma-Aldrich | F1804-50UG | IF 1:200 (methanol) |
| Anti-IL32 | R&D Systems | AF3040 | WB 1:1000, IF 1:250 (methanol/saponin/triton) |
| Anti-LC3 | MBL | PM036 | IF 1:500 (saponin) |
| Anti-p62 | MBL | PM045 | WB 1:2000, IF 1:500 (saponin) |
| Anti-RNF213 | Atlas Antibodies | HPA026790 | WB 1:1000, IF 1:1000 (methanol) |
| Anti-TurboID | Agrisera | AS20 4440 | IF 1:500 (methanol) |
| **Secondary antibodies and stains** |  |  |  |
| Goat anti-rabbit HRP | Invitrogen | 65-6120 | WB 1:5000 |
| Donkey anti-goat HRP | Azure Biosystems | AC2149 | WB 1:5000 |
| Bovine anti-goat CF488A | Biotium | 20293 | IF 1:1000 |
| Bovine anti-goat CF568 | Biotium | 20294 | IF 1:1000 |
| Bovine anti-goat CF640R | Biotium | 20297 | IF 1:1000 |
| Donkey anti-goat AF488 | Invitrogen | A11055 | IF 1:1000 |
| Donkey anti-goat AF568 | Invitrogen | A11057 | IF 1:1000 |
| Donkey anti-goat AF647 | Invitrogen | A21447 | IF 1:1000 |
| Donkey anti-rabbit AF488 | Invitrogen | A21206 | IF 1:1000 |
| Donkey anti-rabbit AF568 | Invitrogen | A10042 | IF 1:1000 |
| Donkey anti-rabbit AF647 | Invitrogen | A31573 | IF 1:1000 |
| Donkey anti-mouse AF568 | Invitrogen | A10037 | IF 1:1000 |
| Donkey anti-rat AF488 | Invitrogen | A21208 | IF 1:1000 |
| Hoechst 33258 | Invitrogen | H21491 | R/IF 1:1000 |
| SYTOX green | Thermo Fisher | S7020 | R/IF 1:10,000 |
| Calcofluor white | Biotium | 29067 | IF 1:1000 (triton) |
| **Cell culture** |  |  |  |
| DMEM | Gibco | 11995073 |  |
| DMEM (phenol red free) | Gibco | 31053028 |  |
| DMEM/F-12, GlutaMAX supplement | Gibco | 10565018 |  |
| EpiCM media | ScienCell | 4101 |  |
| OptiMEM | Gibco | 31985070 |  |
| MEM Non-Essential Amino Acids | Gibco | 11140050 |  |
| GlutaMAX | Gibco | 35050061 |  |
| HEPES (1M) | Gibco | 15630-080 |  |
| Fetal Bovine Serum | Omega Scientific | FB-01 |  |
| Cosmic Calf Serum | Cytiva | SH30087.03 |  |
| 1X PBS, pH 7.4 | Gibco | 10010023 |  |
| 1X HBSS | Gibco | 14025092 |  |
| UltraPure H2O | Invitrogen | 10977023 |  |
| 0.25% Trypsin-EDTA | Gibco | 25200114 |  |
| L-Tryptophan | Sigma Aldrich | T0254-5G |  |
| 6 well cell culture plate, clear | Corning | 3516 |  |
| 24 well cell culture plate,<br>clear | Corning | 3524 |  |
| 96 well cell culture plate,<br>clear | Corning | 3596 |  |
| 96 well cell culture plate,<br>black | Corning | 3904 |  |
| 96 well cell culture plate,<br>white | Corning | 3903 |  |
| **Immunofluorescence**<br>**reagents** |  |  |  |
| 12mm #1.5 glass coverslips | Thermo Fisher | 12541001CA |  |
| Bovine Serum Albumin<br>(BSA) | VWR | 0332-100G |  |
| Glycine | VWR | M103-1KG |  |
| Saponin | Amresco | 0163-100G |  |
| Mowiol 4-88 | Sigma Aldrich | 81381-50g |  |
| 1,4-Phenylenediamine<br>dihydrochloride | Sigma Aldrich | 78460-100g |  |
| **Western blot and**<br>**proteomics reagents** |  |  |  |
| Pierce RIPA buffer | Thermo Fisher | 89900 |  |
| Protease inhibitor cocktail | Millipore Sigma | P8340-5ML |  |
| DNase I | NEB | M0303S |  |
| 4x Laemmli Buffer | BioRad | 1610747 |  |
| β-mercaptoethanol | VWR | M131-100ML | |
| Tween-20 | Sigma Aldrich | P1379-500ML |  |
| 4–20% Mini-PROTEAN<br>TGX Stain-Free Gels | BioRad | 4568096 |  |
| Immobilon-FL PVDF Membrane | Millipore | IPFL00010 |  |
| Blotting-grade blocker (nonfat dry milk) | BioRad | 1706404 |  |
| Clarity Western ECL Substrate | BioRad | 1705061 |  |
| Amersham ECL Prime | Cytiva | RPN2105 |  |
| SuperSignal West Femto ECL | Thermo Fisher | 34095 |  |
| Restore Western Blot Stripping Buffer | Thermo Scientific | 21059 |  |
| Pierce BCA Protein Assay Kits | Thermo Fisher | 89900 |  |
| DTT | Roche | 10197777001 |  |
| MagStrep Strep-Tactin beads | IBA Lifesciences | 2-1613-002 |  |
| DynaMag-2 magnetic separator | Invitrogen | 12321D |  |
| **Cloning** |  |  |  |
| Gateway BP Clonase II Enzyme Mix | Thermo Fisher | 11789020 |  |
| Gateway LR Clonase II Enzyme Mix | Thermo Fisher | 11791020 |  |
| Q5 Site-Directed Mutagenesis Kit | NEB | E0554S |  |
| InFusion HD Cloning Kit | Takara | 639649 |  |
| NucleoSpin Gel and PCR Clean-up Kit | Takara | 740609.50 |  |
| **Assay kits** |  |  |  |
| Human IL-32 DuoSet ELISA | R&D Systems | DY3040-05 |  |
| DuoSet Ancillary Reagent Kit 2 | R&D Systems | DY008B | For IL32 ELISA |
| RealTime-Glo Annexin V Apoptosis and Necrosis Assay | Promega | JA1011 |  |
| Pierce BCA Protein Assay Kit | Thermo Scientific | 23225 |  |
| **Antibiotics, cytokines, inhibitors, transfection reagents** |  |  |  |
| Blasticidin S | Invivogen | ant-bl-1 |  |
| Cycloheximide | Fluka analytical | 1810 |  |
| Epoxomicin | MedChemExpress | HY-13821 |  |
| Geneticin (G418 Sulfate) | Gibco | 10131035 |  |
| Interferon-gamma | Millipore Sigma | IF002 |  |
| Lipofectamine RNAiMAX | Invitrogen | 13778030 |  |
| Lipofectamine 3000 | Invitrogen | L3000008 |  |
| Penicillin-Streptomycin (10,000 U/mL) | Gibco | 15140122 |  |
| Puromycin dihydrochloride | Sigma Aldrich | P9620 |  |
| Tannic Acid | Millipore Sigma | 403040-50G | Used at 80uM for A549 cells |
| TNFα | R&D systems | 210-TA-005 | |
| TransIT 293T Transfection Reagent | Mirus Bio | MIR 2700 |  |
| TransIT LT1 Transfection Reagent | Mirus Bio | MIR 2300 |  |
| Z-VAD-FMK | Invivogen | tlrl-vad |  |
| **Misc** |  |  |  |
| 0.45uM sterile syringe filters | Corning | 431225 |  |
| 5uM sterile syringe filters | PALL | 4650 |  |
| 10X PBS, pH 7.4 | Gibco | 70011044 |  |
| 27 G x 1/2 in.<br>PrecisionGlide Needle | BD | 305109 |  |
| Biotin | Sigma Aldrich | B4501-100MG | Used at 50uM |
| Formaldehyde Solution | Sigma Aldrich | 252549-4L | Diluted to 4% in 1X PBS |
| Methanol | Fisher Chemical | A412-4 | Kept at -20C for permeabilization, room temperature for blocking PVDF |
| Polybrene | Sigma Aldrich | TR-1003-G |  |
| Triton X-100 | ProPure | M143-1L |  |
| TrueCut Cas9 Protein v2 | Invitrogen | A36498 |  |

## PRIMERS

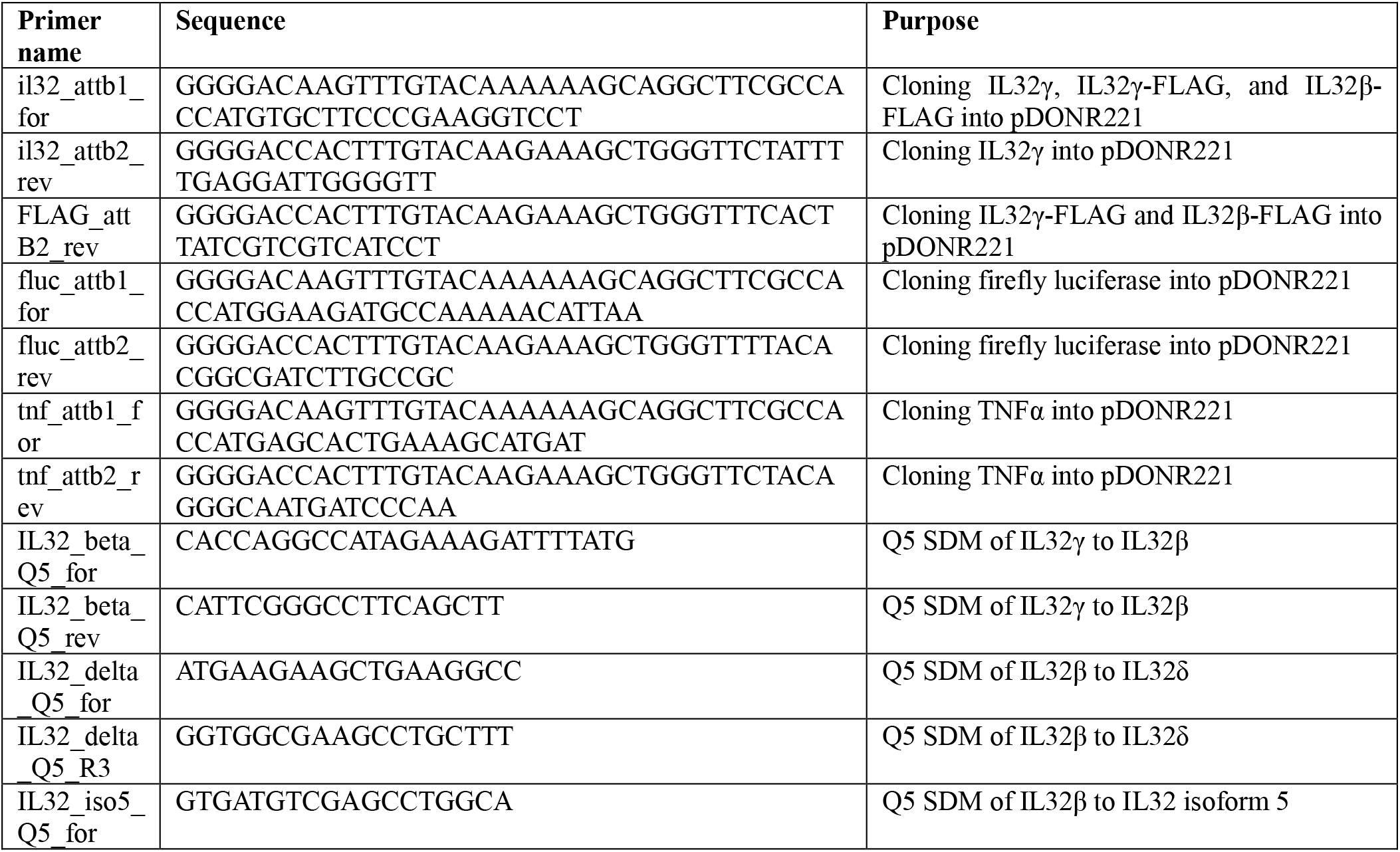
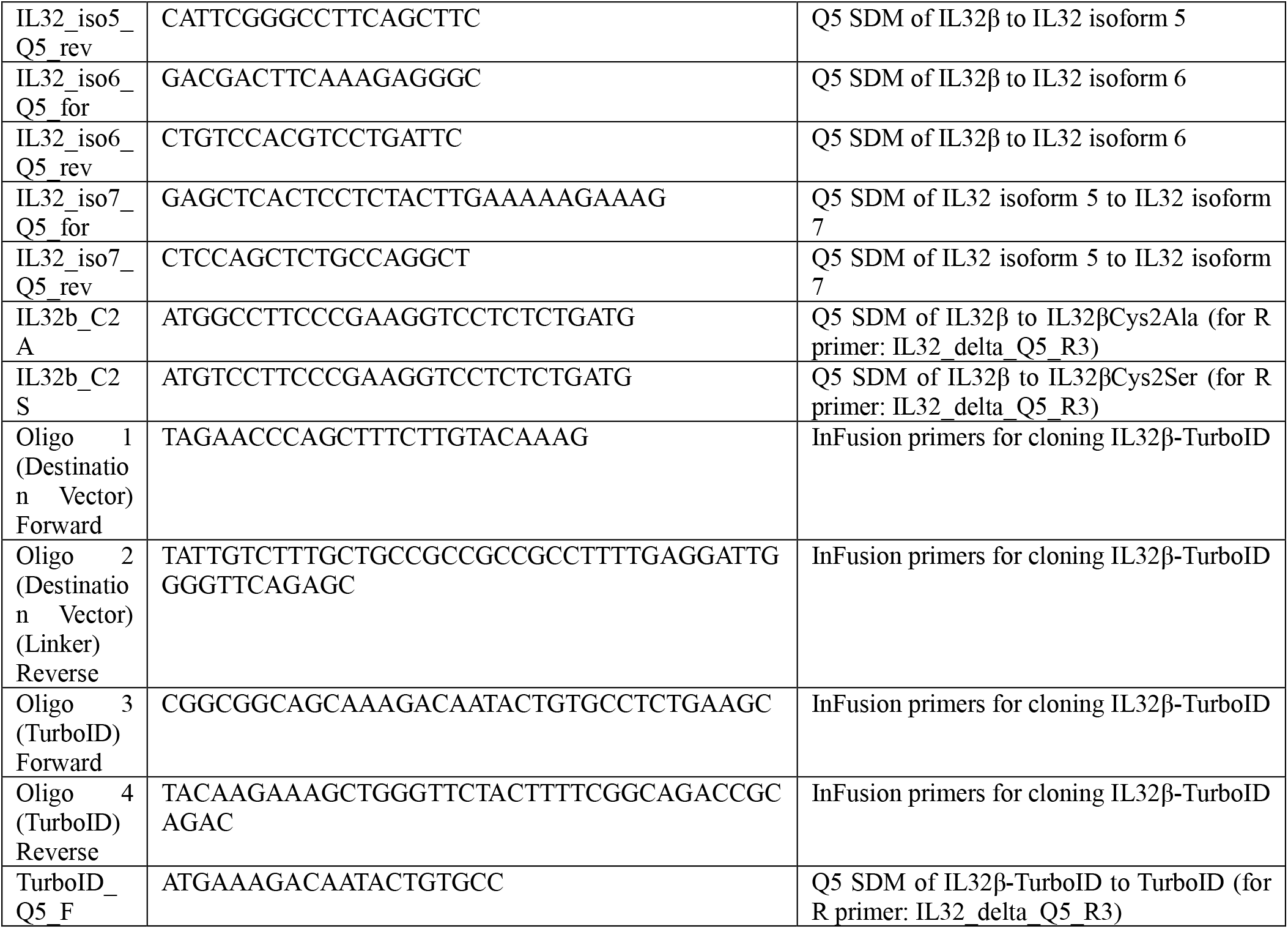

## Acknowledgements

The authors acknowledge the Duke Functional Genomics Shared Resource for their assistance with the execution and analysis of the ISG library and CRISPR screens. We thank the Duke University School of Medicine for the use of the Proteomics and Metabolomics Core Facility and the Duke University Light Microscopy Core Facility (LMCF).

## Author contributions

J.R.R., S.C.W, and J.C. conceived the study and designed the experiments. J.R.R., S.C.W, M.S.D., and E.A. J. performed the laboratory experiments and collected the data. M.E.C., H.A., N.M.A., I.D., K.H., R.S. generated and shared critical reagents. J.R.R., S.C.W, M.S.D., E.A. J., SY.K. and J.C analyzed the data. J.R.R., and J.C. interpreted results and wrote the manuscript. All authors provided critical feedback and reviewed the final manuscript.

